# Mosaic pattern: lung functional heterogeneity at the alveolus level

**DOI:** 10.1101/2025.10.13.682072

**Authors:** Gabrielle N. Grifno, Han Ali Kahvecioglu, Robert LeBourdais, Victoria Travnik, Rohin Banerji, Winita Wangsrikhun, Linzheng Shi, Suleyman B. Bozal, Heewon Suh, Ahmed A. Raslan, Athanasios Batgidis, Byungjun Kang, Feiyang Deng, Caleb Dalton, Andrew Tsao, Lauren Castle, Kathryn Regan, Abdulrahman Kobayter, Michael Vannini, Mohammad Rashidian, Liang Hao, Giovanni Ligresti, Joseph P. Mizgerd, Worth Longest, Michael Hindle, W. Mark Saltzman, James P. Butler, Béla Suki, Hadi T. Nia

## Abstract

Inhaled particles carrying pathogens, pollutants (e.g., microplastics, smoke), therapeutics, and diagnostics are increasingly relevant to public health, yet real-time tracking of aerosol transport in functional alveoli remains challenging. Here, we used the recently developed crystal ribcage to investigate aerosol transport in ex vivo lungs during active ventilation, obtaining the first real-time observations of single aerosol droplet transport and deposition in functional alveoli. We discovered deterministic heterogeneity at both intra- and inter-alveolar levels, with aerosol distribution forming a characteristic "mosaic" pattern in which only specific alveolar clusters received particles. The pattern was consistently formed in vivo during spontaneous breathing and ex vivo using both positive- and negative-pressure ventilation. This pattern was also consistent across a range of aerosols, including small molecules, nanobodies, nanoparticles, microplastics, therapeutics, and pathogens. Additionally, the pattern was observed in murine, porcine, and human lungs, and evolved from birth through aging in mice. The post-deposition stability of the pattern depended on particle type and lung age, lasting from a few minutes for small molecular weight particles to multiple days for cell-binding particles. These alveolar-level heterogeneities may uncover previously unrecognized biological and immunological heterogeneities associated with the mosaic pattern, including its role in postnatal lung development, susceptibility to inhaled airborne hazards such as pollutants and infectious agents, and early pathogenesis and response to inhaled therapeutics in respiratory diseases such as pneumonia, COPD, asthma, and lung cancer.

---

Understanding the inhalation dynamics of microparticles, including pathogenic, therapeutic, and environmental particles, has become increasingly important over the last decade. Airborne pathogens^1,2^, such as viruses and bacteria, are often transported in respiratory droplets. Pollutant particles have raised concerns in urban and occupational settings, with growing attention to microplastics^3^ and particles from household and wildfire smoke^4,5^. Inhalation-based therapeutic^6^, diagnostic^7,8^, and vaccination^9,10^ approaches offer advantages over systemic therapies by bypassing systemic circulation and providing targeted effects^11,12^. Consequently, there is a critical need to study the transport of airborne particles at cellular scales within alveoli, the functional units of the respiratory system. These studies require sufficiently high spatiotemporal resolution to capture the complex dynamics of the interactions between the air and the distending, moving alveolar structures. Achieving real-time, single-particle resolution imaging of airborne particles within a functioning lung, however, remains a considerable challenge.

Animal models have been the gold standard for most inhalation-based studies. However, imaging modalities such as CT and MRI^13-15^ lack the spatial and temporal resolution needed to probe dynamic events such as alveolar deformation, capillary flow, and cellular activities at physiological lung breathing rates. Histological analyses can yield cellular information, particularly with the recent advances in tissue clearing^16,17^ and AI-based 3D pathology^18^. However, these techniques provide static “snapshots” of the tissue without the temporal information associated with aerosol transport into alveoli and their dynamic interactions with alveolar, vascular, and immune cells. Recent intravital microscopy techniques^19-27^ enabled high-resolution imaging of microphysiology dynamics on the lung surface. Despite leading to key discoveries in lung immunology^23,28^ and metastasis^20,29^, these approaches immobilize the lung against the imaging window using either glue or suction, consequently compromising normal respiration mechanics in the area of interest, and they are not compatible with negative-pressure ventilation as in spontaneous breathing^19-21,24-26,28^. Furthermore, intravital microscopy is confined to small regions of the lung, failing to capture whole lobe-scale lung heterogeneities. Lastly, microfluidic systems^30-33^, such as lung-on-a-chip models, can visualize cellular-level events and overcome some limitations of animal studies. However, these models do not recapitulate the complex 3D architecture, heterogeneity, and cellular diversity of the airway and alveolar network, which are essential for studying inhalation dynamics. In summary, despite the microscopic nature of inhaled particles (0.1–10 µm), the alveoli and small airways (50–100 µm), and the affected cells (∼10 µm), current tools and model systems lack the capability to probe airborne particle transport within the native 3D architecture of individual alveoli. Therefore, insights into alveolar-scale aerosol transport have been limited to mathematical modeling^34,35^, lacking direct experimental validation. Consequently, our current understanding of the dynamic interactions between inhaled pathogens, pollutants, and therapeutics at the microscopic level remains incomplete, leaving a wealth of critical phenomena relevant to public health undiscovered.

To address this technological gap, we recently developed the crystal ribcage technology that enables real-time, high-resolution imaging of the whole lung surface while preserving and controlling all three lung phases—gas, liquid, and solid—at the scale of aerosols and cellular components during physiological breathing^36^. By combining controllability and imaging capabilities with the native cellular diversity and architecture of the lung, the crystal ribcage is ideal for visualizing dynamic transport and deposition in the pulmonary (air) phase as well as cellular-level tissue response. Here, we report the first real-time aerosol transport from the whole lobe down to single particles in alveoli during spontaneous breathing as well as negative- and positive-pressure ventilations enabled by crystal ribcage^37^. We discovered stark heterogeneity in aerosol deposition in the lung at the alveolar level, where hotspots of deposition form a deterministic “mosaic pattern” that has never been previously described. The mosaic pattern occurs in alveoli near the pleural surface and throughout the tissue depth and persists in mice and large animals, with indication in human lungs. We probed the evolution of the mosaic pattern across the mouse lifespan and its persistence across a wide range of types and sizes of aerosols relevant to public health, including small molecules, nanoparticles, nanobodies, microplastics, and whole pathogens. We also report how the presence of disease features, such as emphysema and metastatic cancer nodules, remodel the mosaic pattern. The fundamental discoveries presented in this manuscript open new avenues for developing and delivering aerosol-based therapies and diagnostics while providing insight into the links between heterogeneous exposure of alveoli to inhaled materials and the earliest biological and immunological responses to airborne hazards.

## Results and Discussion

### Mosaic pattern is hierarchical and deterministic

To probe the spatiotemporal dynamics of particle transport during breathing down to the alveolar scale, we delivered fluorescent dye dissolved in saline as an aerosol to adult mouse lungs (2-4 months) housed inside the crystal ribcage during continuous ventilation using a modified commercially available nebulizer (**Fig. 1a, Supplemental Video 1**). Surprisingly, aerosols inhaled during lung ventilation did not deposit uniformly across alveoli. Instead, droplets deposited in a spatially heterogeneous but deterministic pattern, where certain alveoli in a near rectangular area preferentially received droplets while neighboring alveoli in the perimeter of the rectangle received no droplets (**Fig. 1b, Supplemental Video 2**). We termed this heterogeneity the “mosaic pattern” due to the tile-like arrangement of regions with high aerosol deposition (**Fig. 1a-b**). Further, we termed regions of high deposition “tiles,” and the surrounding areas of negligible deposition “bands.” Tiles are organized into a multiscale hierarchy from the lobe to the alveolar scale and are self-similar across each spatial scale. The largest tiles, visible even by fluorescent stereo microscopy (**Fig. 1b**), are termed “macrotiles” and are separated by the widest bands of zero droplet deposition, “macrobands”. Each macrotile contains subsets of smaller tiles, termed “mesotiles” that are further comprised of the smallest tiles, the “microtiles,” each containing only a few alveoli. Additionally, we observed that band regions are composed of alveoli and are not occupied by vascular or fissure structures (**Fig. 1b**).

**Fig. 1.**
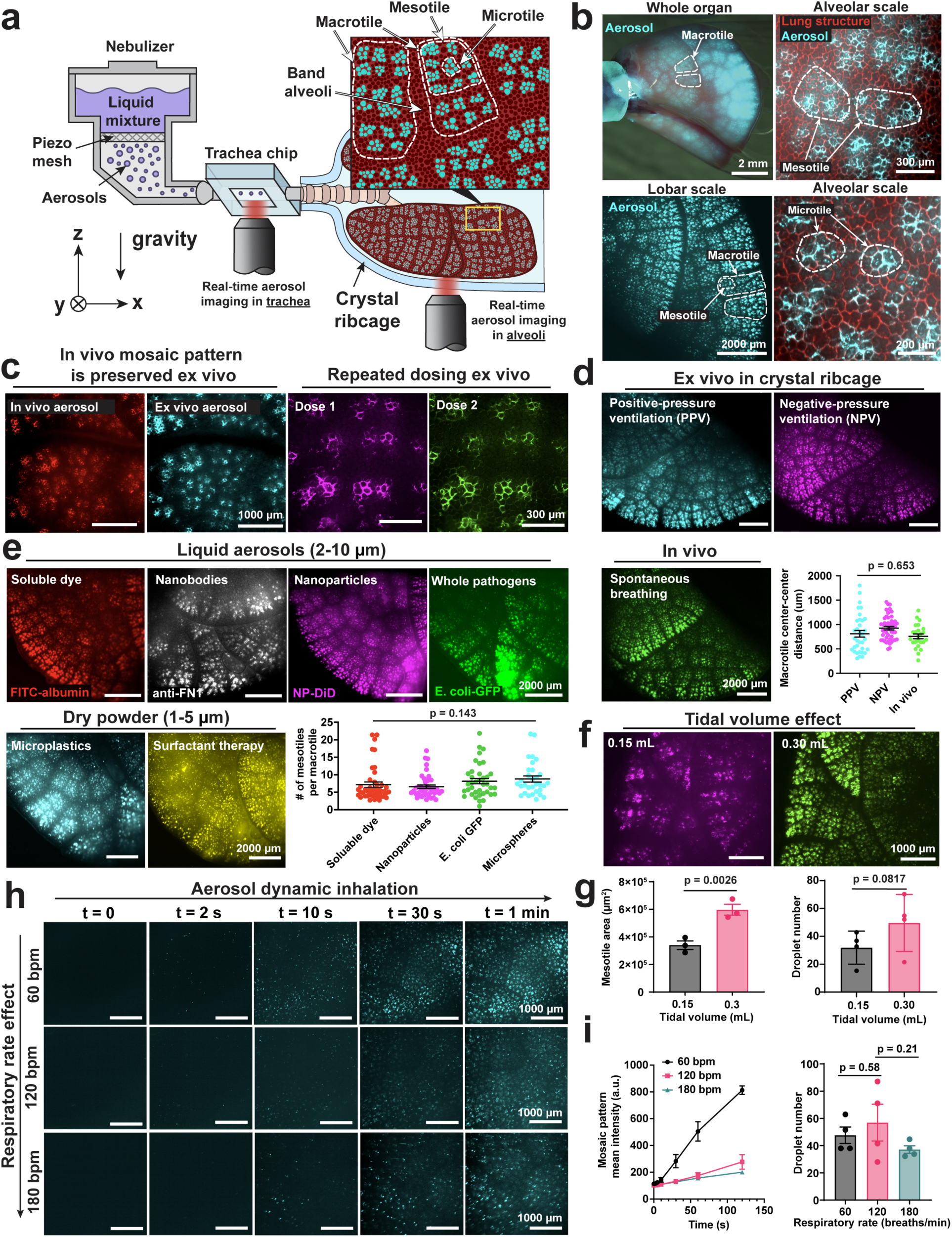
The first observations of the “mosaic pattern” using different aerosolized materials independent of ventilation modality. (**a**) Schematic representation of a whole mouse lung and mosaic pattern formed after aerosol inhalation inside the crystal ribcage, where some populations of alveoli receive droplets while neighboring alveoli do not. (**b**) The mosaic pattern consists of areas of high deposition (bright fluorescence) with a “tile”-like geometry that are surrounded by “bands,” areas of negligible droplet deposition. Tiles are organized into large divisions (macrotiles) with wide spacing and smaller divisions (mesotiles, microtiles) which have closer spacing (n >20 mice). (**c**) The mosaic pattern produced after administering a dose of aerosols in vivo under anesthetized spontaneous breathing and a subsequent dose of aerosols ex vivo by negative pressure ventilation in the same lung are nearly identical, n = 2 independent mouse experiments. Additionally, multiple aerosol doses administered in sequence to the ex vivo lung produce mosaic patterns that are nearly identical (n = 3 independent mouse experiments). (**d**) The mosaic pattern is produced after inhalation of fluorescent aerosols negative- and positive-pressure ventilation and after spontaneous inhalation in anesthetized mice, n = 2-3 independent experiments for each ventilation method, p-value calculated by one-way ANOVA, F = 2.034, df = 51. (**e**) The mosaic pattern is produced using liquid or dry materials with variable properties and characteristic sizes, from droplets (1-10 μm) that contain soluble dyes (1-15 nm), nanobodies (14-18 nm), nanoparticles (100 nm), dry powder synthetic surfactant drug formulations (1 μm), whole pathogens (1 μm), and dry microplastic spheres (1-5 μm). Mosaic pattern heterogeneity, quantified in terms of the number of mesotiles within each macrotile, is conserved across variable materials, n = 1-3 independent mouse experiments, p-value calculated by one-way ANOVA, F = 1.836, df = 156. (**f**) Increasing tidal volume (maintaining respiratory rate (120 breaths/min) and all other parameters consistent) produces a more developed mosaic pattern with larger mesotile structure, n = 3 paired mouse experiments. (**g**) A greater number of single droplets enter the trachea under high tidal volume, m = 4 paired trials across n = 2 independent mouse experiments (mean ± SEM) p-values calculated by two-tailed paired t-test. (**h**) Decreasing respiratory rate results in a faster appearance of the mosaic pattern during dynamic imaging of aerosol inhalation, maintaining tidal volume (0.1 mL) and all other respiratory parameters consistent, n = 3 independent mouse experiments per respiratory rate. (**i**) Mean intensity of the mosaic pattern tiles increases fastest at low respiratory rate, n = 3 independent mouse experiments, while the influx of droplets into the trachea is similar, m = 4 paired trials across n = 2 independent mouse experiments (mean ± SEM), p-values calculated by two-tailed paired t-test.

### Mosaic pattern in spontaneous breathing in vivo vs negative- and positive-pressure ventilations ex vivo

We observed that the hierarchical mosaic pattern is deterministic, not stochastic, and is independent of in vivo vs ex vivo aerosol delivery. We sequentially delivered doses of aerosols using different fluorescent dyes in the same lung, where the first dose (red) was delivered in vivo (spontaneous breathing in anesthetized mice) and the second dose (cyan) ex vivo under ventilation in the crystal ribcage (**Fig. 1c**). The same mosaic pattern was conserved whether aerosols were delivered in vivo or ex vivo within the same lung, as image segmentation revealed that the mosaic pattern tiles formed using each approach were nearly identical (Dice coefficient = 96% ± 0.015% SEM). Additionally, repeated ex vivo doses delivered sequentially but with different fluorescent dyes also retained conserved macro, meso, and microtile structure (**Fig. 1c**, **Extended Data Fig. 1**). This indicates that aerosols are consistently distributed into the same alveoli regardless of whether inhalation occurs via spontaneous breathing or mechanical ventilation.

Furthermore, mosaic pattern formation is also independent of ex vivo ventilation modality. We utilized the key capability of crystal ribcage to observe that the formation of a mosaic pattern in ex vivo lungs using either positive-pressure ventilation (PPV), as in mechanical ventilation, or negative pressure ventilation (NPV) ^37^, are similar to the in vivo mosaic pattern. The periodicity of macrotiles in the mosaic pattern was consistent regardless of ventilation modality (**Fig. 1d**). Additionally, we observed that the brightness of the mosaic pattern is gravity dependent. Single droplets were pulled by gravity toward the lobe of the lung that is the most aligned with the gravity vector (**Extended Data Fig. 2**), causing the mosaic tiles best aligned with gravity to be the brightest (greatest quantity of aerosol deposition).

### Mosaic pattern across environmental, pathogenic, and therapeutic particles

A mosaic pattern consistently formed after inhalation of aerosols containing a wide range of environmental, pathogenic, and therapeutic particles. These materials had variable characteristic size and composition, including nebulized liquid droplets (1-10 µm) containing soluble fluorescent dyes (0.5-70 kDa, approximately 1-15 nm hydrodynamic diameter), nanobodies (camelid heavy-chain variable domains specific to fibronectin, 14-18 kDa^38^), fluorescent nanoparticles (polylactic acid block–hyperbranched polyglycerol, PLA-HPG, ∼100 nm)^39^, whole bacteria *(Escherichia coli*, 1 µm), dry powders comprised of solid microspheres (fluorescent amino formaldehyde polymer, 1-5 µm), and spray-dried powered aerosol formulations of synthetic lung surfactant therapeutics ^40,41^ (**Fig. 1e, Extended Data Fig. 3**). The mosaic patterns had a consistent structure with a similar distribution of mesotiles within each macrotile (**Fig. 1e**) and patterns were formed whether the aerosols were composed of liquid droplets or dry solid particles (**Extended Data Fig. 3**). This implies that inhaled materials, whether harmful or therapeutic, routinely make first contact with the same sub-population of alveolar epithelial and immune cells. As a result, these cells may become particularly tuned to respond to stimuli from pathogens or pollutants through repeated exposure.

Next, we investigated how the mosaic pattern is influenced by key ventilation parameters, tidal volume and respiratory rate, which govern net airflow into the lung. To isolate the effect of tidal volume, we sequentially delivered aerosols in the same lung, where the first dose was delivered under low tidal volume (0.15 mL, 6 mL/kg) and then another dose at high tidal volume (0.3 mL, 12 mL/kg) using two different fluorescent dyes, holding other parameters constant (**Fig. 1f**). This enabled direct visualization of the effect of tidal volume on mosaic pattern heterogeneity. We found that increasing tidal volume led to a more developed mosaic pattern across spatial scales, leading to an increase in macrotile and mesotile area (**Fig. 1f-g, Extended Data Fig. 4**). To understand this result, we imaged the influx of single droplets using a tracheal imaging microfluidic chip (**Fig. 1a**). The results revealed a trend that higher tidal volume leads to a greater number of droplets entering the lung (**Fig. 1g**). We found that increasing tidal volume increases the influx of single droplets into the trachea, meaning that more droplets eventually transit to the distal alveoli and deposit on the lung surface, resulting in a consistent increase in tile size at multiple scales of the mosaic pattern.

We next probed how respiratory rate affects the rate of mosaic pattern formation. Using ultrafast spinning disk microscopy during continuous ventilation, we imaged real-time mosaic pattern formation under different respiratory rates (60, 120, and 180 breaths/min), keeping all other parameters constant (**Fig. 1h**). Decreasing breathing rate led to faster mosaic formation, despite similar particle influx into the trachea across rates (**Fig. 1h-i**, m = 4 paired trials, n = 2 mice) measured via the trachea chip. Additionally, changes in respiratory rate did not affect mosaic tile size, across multiple spatial scales (**Extended Data Fig. 5**). This suggests that, instead of modulating total particle inspiration, respiratory rate alters fluid mechanics in the deeper airways and alveoli. The Stokes number (Stk), which characterizes a particle’s ability to follow fluid motion, is defined as:

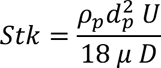

Where *ρ*_*p*_ is the density of the particle, *d*_*p*_ is the diameter of the particle, *U* is the fluid velocity, *μ* is the fluid dynamic viscosity, and *D* is the characteristic length of the system. Increasing respiratory rate, while keeping the tidal volume constant, leads to proportional increase in *U* and hence Stk number. The larger the Stk, the more particles move independently from the flow, following their inertia rather than the fluid streamline. Thus, due to inertial impaction, higher respiratory rates increase aerosol deposition in the large airways and at airway bifurcation points, such that fewer particles transit to the distal alveoli. This is consistent with our observation that at a higher respiratory rate longer times are needed to see enough particle accumulation to form a visible mosaic pattern on the lung surface.

### Mosaic pattern evolves across the lung lifespan and persists across species

As lung development continues after birth^42^, with rapid alveologenesis occurring 4-7 days post birth in mice^43,44^, we asked whether the mosaic pattern is present from birth or emerges due to postnatal development of alveoli. We delivered liquid aerosols containing cell membrane-binding dye to neonatal mouse lungs under anesthetized spontaneous breathing on days 0, 3, 4, and 7 after birth and compared the resulting aerosol distribution pattern to adult mouse lungs (2-4 months) and aged lungs (20-28 months). Interestingly, we observed that the spatial organization of the mosaic pattern is conserved from birth throughout the lifetime of the lung (**Fig. 2a**). However, the number of alveoli contained within each tile increases steadily with age, while the bands between tiles are consistently composed of only 1-3 alveoli across the lifespan of the mouse (**Fig. 2b**). This may indicate that postnatal alveologenesis may be associated with the mosaic pattern with potentially important biological consequences for future studies in lung development. Additionally, as alveoli in the tiles have greater access to inhaled particles, they may also have greater access to gases. Differential exposure may mean that alveoli in tiles receive differential mechanical and biological cues that contribute to postnatal lung development.

**Fig. 2.**
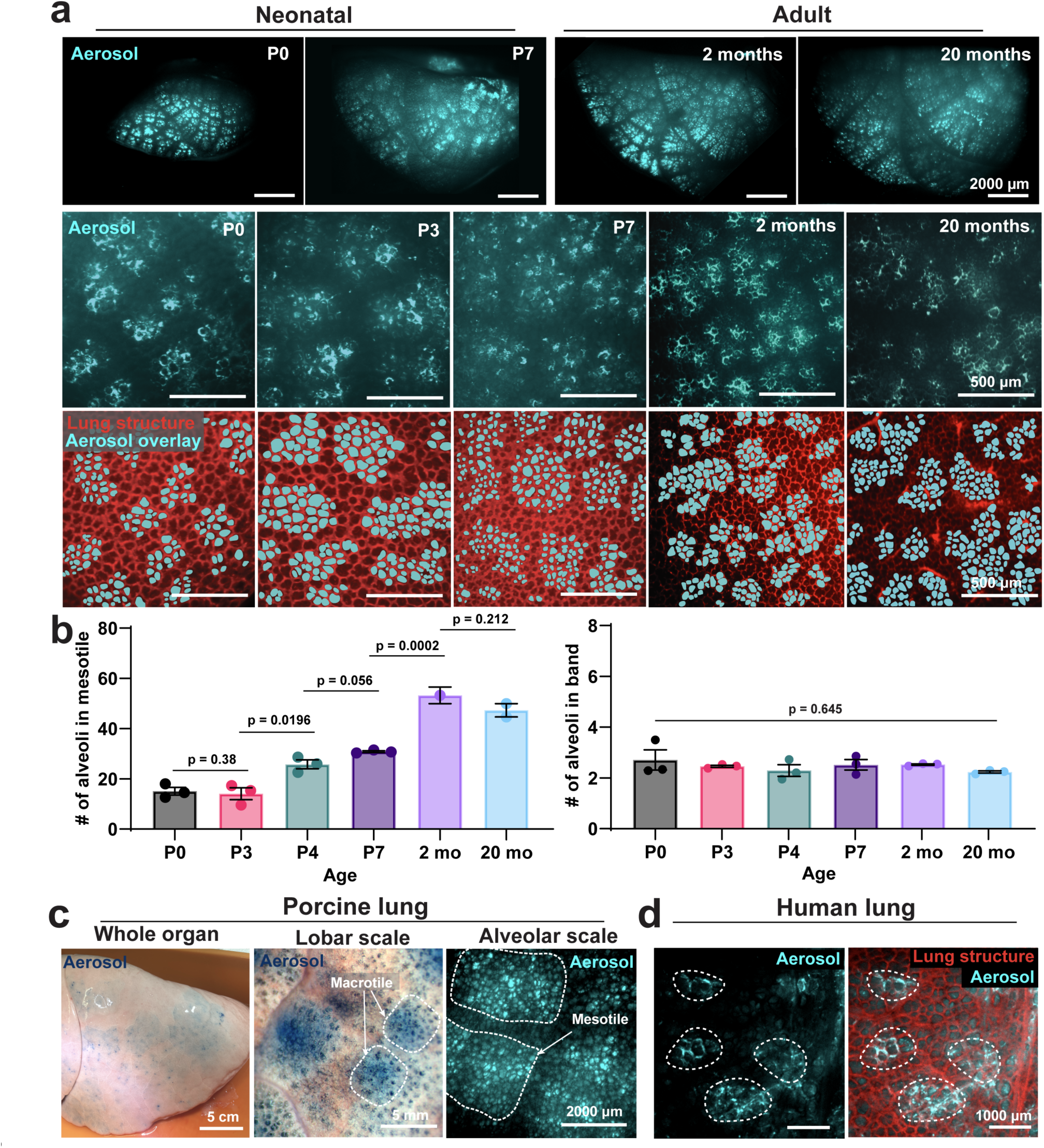
The mosaic pattern evolves across the mouse lifespan and is present across species. (**a**) The mosaic pattern is seen after aerosol inhalation across the whole lifespan of the mouse, from postnatal day 0 to two years, n = 3 mice per age group. (**b**) The number of alveoli in the width of the mesotiles increases with age (mean ± SEM, p-values calculated by Welch’s t-test), however, the number of alveoli in the width of the band between tiles remains constant with age (mean ± SEM, p-value for all ages calculated by one-way ANOVA, F = 0.6831, df = 17). (**c**) A mosaic-like pattern is present on the surface of porcine lungs after aerosol inhalation (n = 2 independent pig lungs), revealed by stereomicroscopy and spinning disk confocal microscopy. (**d**) Indication of mosaic-like heterogeneity of aerosol deposition seen in human lungs (n = 2 independent human lungs), obtained by spinning disk confocal microscopy.

To confirm that the mosaic pattern is not limited to mouse lungs and is relevant to lungs with similar size and architecture to humans, we demonstrated that the mosaic pattern is also present in porcine lungs. After aerosol delivery to adult porcine lungs, a mosaic-like pattern of deposition appeared on the pleural surface, indicating that this heterogeneity persists in large animals with lung sizes comparable to humans and is not unique to mice (**Fig. 2c, Extended Data Fig. 6**). Finally, we investigated the presence of the mosaic pattern in human transplant-rejected lungs. Despite the non-ideal condition of transplant-rejected lungs and multi-hour sample transport postmortem, we also observed mosaic-like heterogeneity in aerosol deposition in human lungs after delivery of aerosols during ex vivo ventilation (**Fig. 2d, Extended Data Fig. 7**). These observations suggest that the mosaic pattern, with implications for development and disease, is likely a fundamental aspect of normal physiology in large animals and humans.

### The origin of the mosaic pattern

The presence of the mosaic pattern throughout the lung lifespan and its deterministic nature suggest that the pattern arises from fundamental attributes of the lung architecture and breathing mechanics. Dynamic imaging of single droplet arrival in alveoli shows that the mosaic pattern initially forms by direct droplet deposition rather than post-deposition diffusion. After several seconds of single droplet accumulation only in tile alveoli, the overall pattern structure begins to emerge (**Extended Data Fig. 8, Supplemental Video 3**). To probe the dynamic mechanisms that are responsible for mosaic heterogeneity, we first asked whether the pattern is associated with alveolar mechanics and whether it can be modulated by perturbing lung airway structure or alveolar structure. Thus, we hypothesized that local differences in alveolar expansion could explain the difference in deposition between adjacent tile and band alveoli. However, examining alveolar area as a function of inflation pressure and the areal strain^45^ in alveoli, we found that strain is similar between the tile and band (**Extended Data Fig. 9a**). Aerosol deposition also did not affect alveolar area in response to dynamic ventilation after aerosol delivery **(Extended Data Fig. 9b-c**). Additionally, we probed whether the mosaic pattern heterogeneity is related to respiration-circulation coupling. After labeling the mosaic pattern in vivo, we perfused a bolus of dye through the vessels in the ex vivo lung inside the crystal ribcage but observed no association between the mosaic pattern and perfusion through the pulmonary capillaries (**Extended Data Fig. 10, Supplemental Video 4**).

After demonstrating that alveolar and vascular mechanics are not associated with the mosaic pattern, we investigated whether the mosaic pattern arises from airway structure. Imaging precision-cut lung slices (PCLS) of the left lobe after aerosol delivery revealed that the mosaic pattern is not a localized phenomenon at the pleural surface but also exists deeper in the parenchyma (**Fig. 3a**). Tiles with high aerosol deposition were connected to lung terminal bronchioles, while bands occupied the space between airway branches after bifurcations. This suggests that aerosols preferentially travel along the straightest (and likely least resistant) paths before depositing in alveoli, as similar flow distributions have been previously suggested^46^. When the paths become narrow and tortuous farther from the bronchioles and alveolar ducts, aerosol deposition drops off, leading to mosaic pattern heterogeneity between neighboring populations of alveoli.

**Fig. 3.**
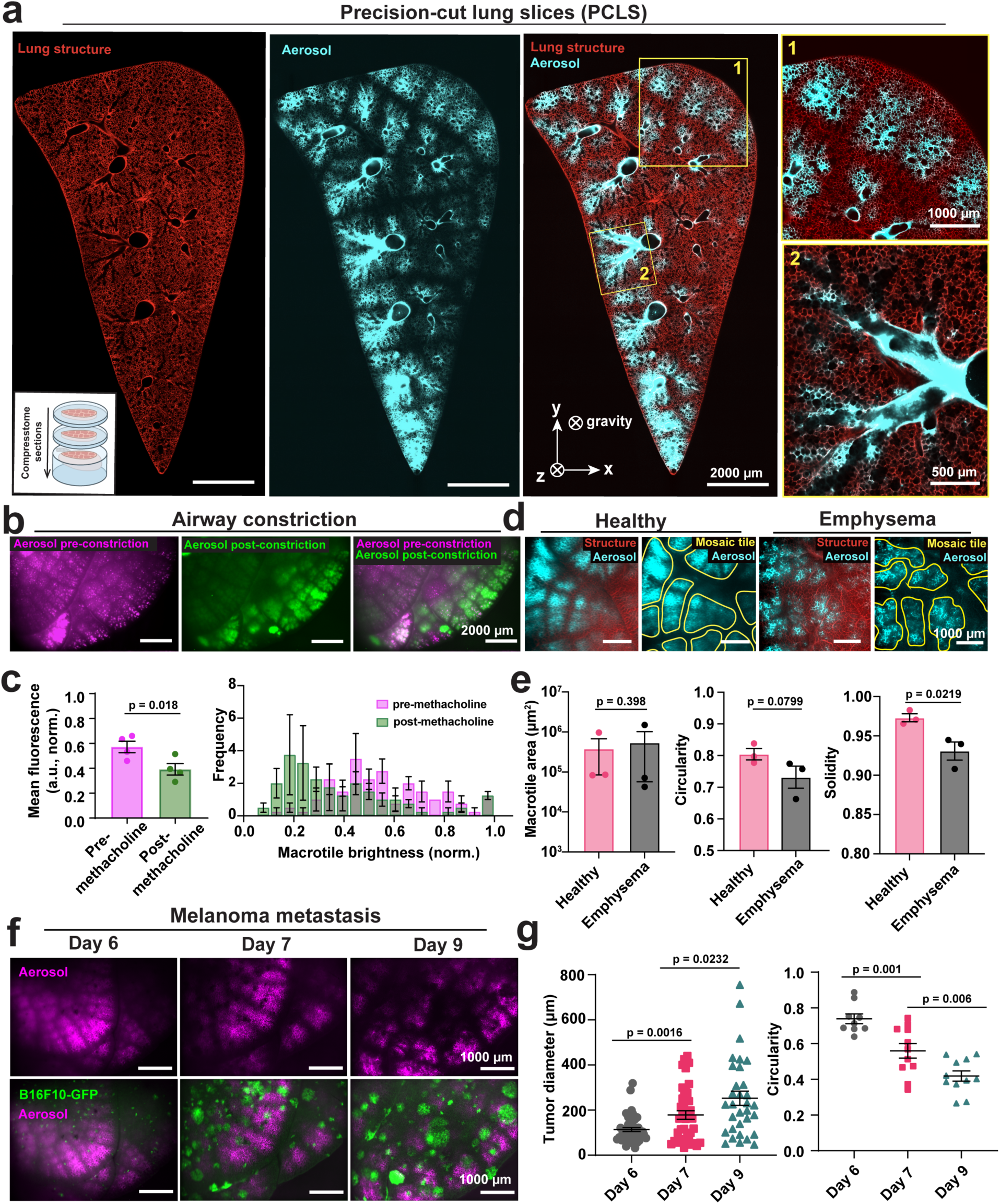
Mosaic pattern heterogeneity is dependent on airway structure. (**a**) The mosaic pattern persists throughout the lung parenchyma, observed via spinning disk confocal images of precision-cut lung slices; representative image located 4 mm below the pleural surface. Tiles where aerosols preferentially deposit are spatially closer to terminal airways, n = 3 independent mouse experiments. (**b**) Aerosol inhalation before and after airway constriction achieved by methacholine challenge was used to directly probe the mosaic pattern as a function of airway functionality. (**c**) Airway constriction caused tile mean fluorescence to decrease compared to the mosaic pattern prior to constriction, n = 4 independent mouse experiments (mean ± SEM), p-value calculated by Welch’s t-test. (**d**) Lung alveolar structure is heavily remodeled in an elastase model of emphysema. (**e**) Tiles cover similar spatial area on the lung surface in health and emphysema, but injured tiles have irregular geometry compared to healthy tiles, n = 3 mouse experiments (mean ± SEM), p-value calculated by Welch’s t-test. (**f**) Melanoma metastasis to the lungs causes a solid phase obstruction in the local alveolar airspaces. However, tumors do not heavily remodel the mosaic pattern when obstructions are small, but disruption increases as tumors progress (n = 1-2 independent mouse experiment per timepoint). (**g**) Tumor diameter and mosaic macrotile circularity quantified for 32-45 tumors and 9-11 mosaic tiles per mouse, n = 1 independent mouse experiment per timepoint (mean ± SEM), p-values calculated by Welch’s t-test.

To test the hypothesis that airway structure is the dominant driver of the mosaic pattern, we delivered aerosols to the lung before and after intravascular delivery of methacholine (**Fig. 3b-c, Extended Data Fig. 11**). Methacholine constricts lung airways by causing contraction of smooth muscle cells^47^. We delivered methacholine intravascularly to ensure uniform distribution through the lung tissue. Additionally, intravascular delivery ensured that airways would be free of excess liquid during aerosol delivery, which could hinder aerosol transport especially to the distal alveoli. We delivered a dose of fluorescent aerosol droplets before and after methacholine challenge to probe how flow through the airways contributes to the mosaic pattern. We compared the “normal” mosaic pattern without perturbation to the pattern formed after constriction. Many of the tiles that initially had high aerosol deposition (and thus high fluorescence intensity) prior to constriction had reduced intensity after airway constriction, although the overall pattern structure was conserved (**Fig. 3c**). Additionally, after the administration of methacholine, the mean fluorescence across all tiles decreased (**Fig. 3c**). These results indicate that the spatial organization of the mosaic pattern is intrinsic to the lung airway structure, as the locations of the tiles do not change with airway constriction. However, the changes in tile brightness that occur after methacholine illustrate how airway constriction reduces the amount of flow through some airways, resulting in reduced droplet deposition (and thus fluorescence intensity) in some tiles.

To determine whether widespread structural changes at the alveolar level can overturn the dominance of airway structure in producing the mosaic pattern, we probed pattern formation in mouse models of emphysema, fibrosis, and metastasis. Elastase delivered to healthy lungs causes rupturing of the septal wall between alveoli, creating large gaps in the tissue similar to emphysema in humans^48,49^ (**Extended Data Fig. 12**). However, despite drastic remodeling of the alveolar structure, the mosaic pattern was still present with a similar tile size distribution and organization (**Fig. 3d**). Tiles covered approximately the same spatial area on the lung surface but had more irregular geometry (decreased circularity and solidity) compared to tiles in healthy lungs (**Fig. 3e**). In a mouse model of pulmonary fibrosis, which affects both airways and alveoli, tissue remodeling and increased collagen in heavily injured regions prevented the formation of the mosaic pattern, while the pattern appeared normally in less injured regions (**Extended Data Fig. 13**). These trends indicate that alveolar-level structural changes are unlikely to dictate major changes in mosaic pattern as long as airflow is present. Lastly, solid metastasis is known to remodel lung structure and leads to local disruption of alveolar and capillary mechanics^36,50,51^. However, how solid tumors disrupt aerosol transport at the alveolar level is poorly understood. In a mouse model of melanoma metastasis, tumors did not substantially alter the mosaic pattern when the tumor diameter was on the same scale as alveoli (<200 µm, day 6) (**Fig. 3f-g**). Instead, tumors needed to reach a critical size (>200 µm, day 7-9) to disrupt the mosaic pattern tile geometry. Taken together, these results provide evidence that the larger airway structures are dominant in forming the pattern even in the presence of alveolar-scale irregularities in the tissue such as small clusters of cancer cells.

### Mosaic pattern stability is age and molecular weight dependent

We next investigated how the mosaic pattern evolves in the minutes to hours following deposition, assessing whether interstitial transport contributes to delayed exposure in initially unexposed alveoli. Continuous observation of the lung revealed that alveoli in the bands that did not directly receive droplets became exposed to small molecule fluorophores in a secondary redistribution phase (**Fig. 4a**). Soluble fluorescent small molecules and proteins (0.5-70 kDa, 1-15 nm hydrodynamic diameter) dissolved inside liquid droplets redistributed throughout the lung and caused blurring of the mosaic pattern within 10 minutes (**Fig. 4b-c**, **Supplemental Video 5**). Similarly, nanobodies (15 kDa) redistributed quickly over minutes before binding uniformly to lung extracellular matrix structures (**Fig. 4b-c**). Additionally, the rate of mosaic tile blurring decreased with increasing material molecular weight.

**Fig. 4.**
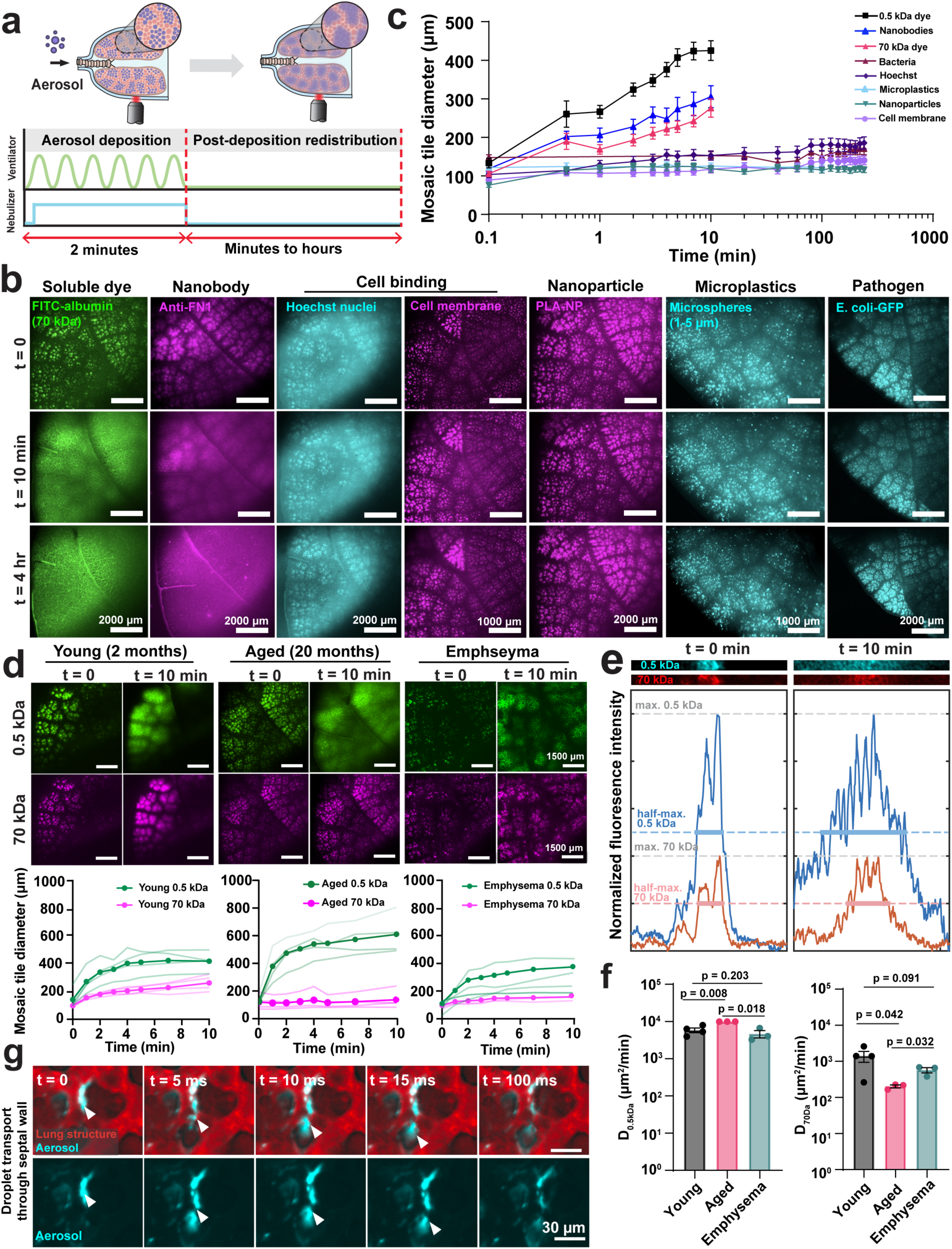
Aerosol redistribution post-deposition is age- and molecular weight-dependent. (**a**) Schematic representation of aerosol redistribution post droplet deposition in the absence of ventilation after aerosol inhalation. (**b**) Soluble small molecules and nanobodies diffuse over 10 min; nucleus (Hoechst 33342) or cell membrane (DiD)-binding dyes co-delivered to the lung bind to cells upon contact and do not redistribute over 4 hours; nanoparticles (300 nm) and dry microspheres (1-5 µm) do not redistribute over 4 hours due to their size; pathogens do not redistribute in 4 hours due to their large size. (**c**) Quantification of mosaic tile diameter (µm) over time for each material, n = 2-3 independent experiments per material (mean ± SEM), m = 5-10 ROI measured per experiment. (**d**) In young mice, liquid aerosols co-delivering 0.5 kDa (“lightweight”) and 70 kDa (“heavy”) fluorescent tracers redistribute from the mosaic tiles to the surrounding bands after depositing onto the tissue surface at rates proportional to molecular weight (n = 4 mouse experiments, m = 5-10 ROI per experiment). In aged mice, redistribution of heavy fluorophores is slower than in young mice (n = 4 mouse experiments, m = 5-10 ROI per experiment). Elastase injury in a mouse model of emphysema heavily remodels alveolar structure, altering the time dynamics of post-deposition aerosol redistribution after inhalation of a mixture of lightweight and heavyweight dyes (n = 3 mouse experiments, m = 5-10 ROI per experiment). (**e**) Measuring the fluorescence intensity across a line profile drawn over the mosaic mesotiles allows for quantification of the rate of transport for each dye over time. (**f**) The diffusion coefficient of aerosols during redistribution in the lung tissue is measured after fitting the line profile data to a spherical model of diffusion, (n = 3-4 mouse experiments per condition, m = 5-10 ROI per experiment, mean ± SEM, -values calculated using Welsh’s t-test). (**g**) Single droplets that deposit on the alveolar wall spread thinly on the alveolar wall and pass through the septal wall into neighboring alveoli milliseconds after initial impaction (n = 3 independent mouse experiments).

Interestingly, soluble dyes that quickly bind to specific cellular structures such as Hoechst 33342 (nuclei) and DiD (cell membranes) did not substantially redistribute and retained a stable microtile organization upon droplet contact with the tissue. Fluorescent nanoparticles (300 nm diameter) and dry powder microplastic spheres (1-5 μm) also did not redistribute over several hours due to their large particle size. Similarly, whole bacteria (E. coli-GFP, 1-2 μm) did not redistribute due to their large size in the hours following aerosol delivery (**Fig. 4b-c**). Quantification of real time aerosol redistribution at the lobar level will have implications for both drug delivery and disease and injury models in the lung, as material properties determine whether different groups of alveoli become exposed to inhaled materials by direct deposition or later redistribution. Direct insight into the aerosol deposition and clearance (or retention) of these materials is crucial for understanding how immune responses are spatially and temporally organized given that only particular alveoli come into routine direct contact with immunogenic aerosols.

To probe the multiday stability of mosaic pattern, we administered aerosols to mice in vivo, for one example material (cell membrane binding dye) (**Extended Data Fig. 14**). Monitoring the same lung in vivo was not possible, so mice were sacrificed daily for three days for imaging. The mosaic pattern was retained for up to 2 days, after which it dissipated, potentially by uptake by cells such as alveolar macrophages, such that discrete tiles were no longer visible on day 3. The multiday retention of mosaic pattern heterogeneity implies long-term heterogeneous distribution of therapeutics, pollutants, and pathogens that could cause biological differences between alveolar cells in the band versus tiles.

Next, we probed how variations in lung tissue properties associated with aging impact mosaic pattern stability. In young adult mice (2-4 months), liquid aerosols containing both 0.5 kDa fluorescent tracers (modeling small-molecule drugs) and 70 kDa fluorescent tracers (modeling whole proteins) redistribute from the mosaic tiles to the surrounding bands after depositing onto the tissue surface at rates inversely proportional to dye molecular weight (**Fig. 4d**). Measuring the fluorescence intensity across a line profile drawn over each mosaic tile allowed for quantification of the rate of transport for each dye over time (**Fig. 4e**). After fitting this line profile data to a spherical model of diffusion, we quantified the diffusion coefficient, D, for soluble dyes redistributing in mouse lung tissue (**Fig. 4f**). The resulting D values were similar to diffusion coefficients of Lucifer Yellow salts and bovine serum albumin in water reported in the literature, at 3 x 10^4^ µm^2^/min ^52^ and 3.6 x 10^3^ µm^2^/min ^53^, respectively. Additionally, molecular weight-dependent redistribution was conserved across variable materials. We repeated the experiments with fluorescent salts and albumins in Fig. 4 with small (3 kDa) and large (70 kDa) fluorescent dextrans and found that the molecular weight redistribution was much slower for heavier dextrans in young adult mice (2-4 months), similar to the trend seen between small molecules and fluorescent albumins (**Extended Data Fig. 15**).

Interestingly, in aged mice (20-28 months), the rate of heavy fluorophore (70 kDa) redistribution decreased significantly. (**Fig. 4d, Extended Data Fig. 16, Supplemental Video 5**). The diffusion coefficient for 70 kDa dyes was reduced by an order of magnitude in aged lungs (**Fig. 4f**), which suggests that local tissue (epithelial, immune) response to harmful airborne particles may be remodeled in aging. In turn, this has implications for changes in local alveolar mechanics and breakdown of cellular repair mechanisms associated with aging. Interestingly, we found that in emphysema, the rate of large fluorophore redistribution also showed a decreasing trend compared to healthy lungs (**Fig. 4d-f**). Differences in post-deposition aerosol dynamics in young and aged mice (relative to the mosaic pattern) will provide critical insight into early pathogenesis of age-related diseases such as COPD and cancer. Additionally, slower transport dynamics for soluble molecules in aging may mean that certain populations of alveoli are not accessible to inhaled therapeutics, changing our understanding of the efficacy of different therapies that target the lung in young versus elderly patients.

To determine what mechanisms could explain the differences in redistribution between young and aged (or emphysematous) lungs, we probed how soluble dyes are transported from alveolus to alveolus immediately after aerosol deposition. We observed that single droplets that deposit onto the cell membrane spread along the air-liquid interface of the alveolar wall but then pass through the septal wall into a neighboring alveolus within milliseconds after initial impaction. This direct transport between neighboring alveoli likely happens through pores of Kohn, perhaps driven by a difference in surface tension between alveoli on either side of the pore, as the transport is too fast (milliseconds) to be driven by simple diffusion (**Fig. 4g, Supplemental Video 6**). This suggests a new functional role (that may be remodeled in aging) for pores of Kohn in facilitating transport of liquid aerosols, in addition to the recent finding that they provide conduits for cell migration^54^ and their potential role in collateral ventilation^55-57^.

The heterogeneity of the mosaic pattern, and the age- and material-dependent variability in post-deposition transport suggest that exposure of alveolar and interstitial cells (epithelial, endothelial, and resident immune cells) to aerosols is spatially heterogeneous. Routine exposure to inhaled pollutants and pathogens over the lifetime of an organism may locally “tune” alveolar cells to particular signaling pathways. Lung epithelial cells (e.g., type 2 cells or club cells) include discrete subsets with distinct transcriptomes, surface markers, and functions^58-61^, and such lung cell heterogeneities could result from different exposure histories, depending on location within the mosaic pattern. These epithelial cells coordinate signaling and recruitment of immune cells including alveolar macrophages, neutrophils, eosinophils, and lymphocytes^62^, so responses to inhaled challenges may be profoundly influenced by the effects of the mosaic pattern on epithelial cells. Additionally, alveolar macrophages responding to harmful aerosols localized in mosaic tiles may become overwhelmed or activated disproportionately in comparison to cells located in bands. This may lead to ineffective clearance or localized inflammatory damage. Alveolar macrophages may also be phenotypically distinct, particularly if some preferentially patrol tile or band regions. Finally, recovery from infection leads to persistent immune changes that are confined to previously infected lung regions^62^, including diversification of cell phenotypes (such as alveolar macrophages with different transcriptomes and metabolomes) plus the addition of new cells in the tissue (such as resident memory lymphocytes). Lung cell biology mediating immune resistance and tissue resilience may include regional variations that depend on the lung’s history of mosaic exposures, now becoming an important new focus of investigation.

With respect to therapeutics, heterogeneous transport of aerosols also suggests that alveoli in mosaic “bands” likely experience delayed or reduced exposure to inhaled therapeutic drugs larger than a typical protein. To improve drug delivery in these regions, therapeutic strategies could include development of drugs that can easily diffuse along the air-liquid interface by or repeated dosing regimens to increase exposure of the band alveoli to therapies. In lung aging, some alveoli may be entirely inaccessible to large-molecule or nanoparticle-sized therapies, which would be a key factor to consider when developing inhaled treatments for elderly patients. Similarly, if immune responses are spatially dependent on the mosaic pattern, inhaled therapies may be more effective than systemic administration to elicit an immune response to infections. Lastly, modulation of the mosaic pattern in diseases such as emphysema, fibrosis, and metastasis could indicate how inhalable medicines can be designed to target different diseases and be modulated according to the stage of disease progression.

### Intra-alveolus heterogeneities of single aerosol droplet transport

The *inter*-alveolar heterogeneity of the mosaic pattern prompted us to next leverage the crystal ribcage to investigate whether aerosol transport was also heterogeneous at the *intra*-alveolar scale, which would reveal additional insight into the physics of droplet transport. Coupling the crystal ribcage with ultrafast spinning disk confocal microscopy has enabled the first real-time visualization of single aerosol droplet arrival and deposition on the interior alveolar surface during ventilation. We imaged single aerosols both through a transparent tracheal chip fixed between the mouse lung and nebulizer and in the alveoli through the crystal ribcage. From this microscopy data, we measured the size distribution of aerosols (1-10 μm) during active inhalation as droplets entered the trachea and after transit to the distal alveoli, respectively (**Fig. 5a-b, Extended Data Fig. 17, Supplemental Video 7**). We validated the optical measurements of single droplet diameter using a particle size analyzer and found that optical measurements had a percentage error of 12.4%, which we believe is acceptable for the conclusions in this study (**Extended Data Fig. 18)**. Particles were within a 1-10 μm range using both optical and laser diffraction measurement techniques for various materials and across different nebulization devices (**Extended Data Figs. 18-19**). Aerosols measured by optical microscopy in the tracheal chip had a mean size of 5.5 μm (SEM = 0.063 μm) in comparison to 4.1 μm (SEM = 0.173 μm) inside the alveolar airspace (**Fig. 5b, Supplemental Video 7**). The number of droplets observed in the airspace peaked at approximately 5 droplets in the field of view after peak inhalation. However, replacing the gas medium with helium resulted in an order of magnitude increase in the number of droplets observed at nearly all points during the ventilation cycle (**Fig. 5b**).

**Fig. 5.**
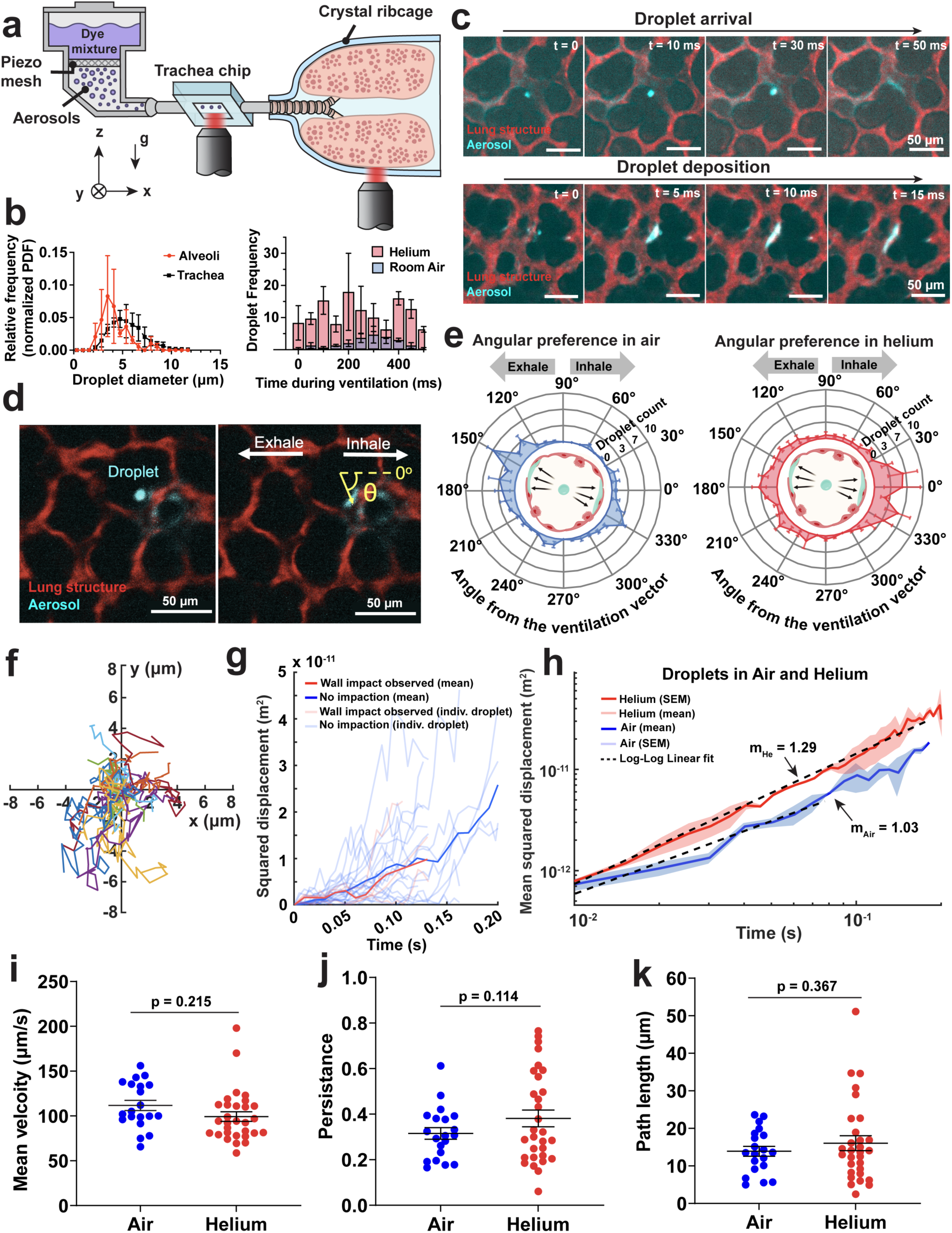
Single aerosol transport is heterogeneous in the pulmonary (air) phase. (**a**) Schematic representation of the system used to image aerosols during inhalation at the mouse trachea and in alveoli. (**b**) Size distribution of aerosols at the trachea versus in alveoli (normalized PDF, where probability is equal to the histogram bin counts divided by number of elements, divided by the bin width, n = 2 paired experiments, mean ± SEM), and increase in aerosol droplets observed at the alveolar level in helium as compared to room air (n = 2 experiments, mean ± SEM). (**c**) Timelapse images of droplet arrival into the alveolar airspace and deposition on the alveolus air-liquid interface during dynamic ventilation using an ultrafast spinning-disk microscope (n = 4 independent mouse experiments). (**d**) Schematic representation of droplet angle of deposition measurement. (**e**) Droplets deposit onto the alveolar wall with an angular preference for the sides of the alveolus aligned with the rigid body motion of the lung during active ventilation, n = 3-4 mouse experiments (mean ± SEM). (**f**) Trajectory map of single droplet paths traveling through room air in the absence of ventilation, n = 2 independent mouse experiments, m = 20 independent droplets tracked. (**g**) Squared displacement of individual single droplets trajectories computed for droplets that impact on the alveolar wall (red) and droplets that leave the focal plane before impaction (blue), n = 2 independent experiments, m = 20 independent droplets tracked. (**h**) Log-log plot of the mean squared displacement, where slope corresponds to the diffusivity of droplets in room air (blue) and helium (red), respectively, with linear fit (dashed black line), n = 2 independent experiments, m = 20 independent droplets per gas composition, (**i**) mean velocity, (**j**) path persistence (total displacement divided by path contour length), and (**k**) total path length measured for individual droplets trajectories and are similar regardless of gas medium used, (n = 2 independent experiments, m = 20-30 independent droplets tracked, mean ± SEM), p-values calculated by Welch’s t-test.

Upon arriving in alveoli, aerosols remained in the air for 30–50 ms before depositing onto the alveolar wall. Once droplets contacted the wall, they spread along the thin layer of surfactant on the surface in the following 15-30 ms (**Fig. 5c, Supplemental Video 8**). In the lung, we observed that aerosol deposition was spatially heterogeneous even within individual alveoli during ventilation, challenging the common assumption that they should impact the alveolar wall with equal probability, due to transport being dominated by random-walk diffusion at the alveolar level. Instead, droplets preferentially deposited on the alveolar wall in locations that were aligned with the direction of the rigid body motion of the lung alveoli during breathing (**Fig. 5d-e, Supplemental Video 9**). This angular preference indicates that aerosol deposition is not just directed by diffusion, as was previously predicted by computational modeling of aerosol transport^34,35^. Additionally, alveoli deformed isotropically (deformation was equal in all angles in the X-Y plane) during ventilation inside the crystal ribcage (**Extended Data Fig. 20**), indicating that alveolar deformation is not responsible for heterogeneous intra-alveolar distribution.

Utilizing the capabilities of the crystal ribcage, we repeated this experiment with both air and helium as the inhaled gases to probe the effect of gas density on single droplet transport dynamics. To investigate the motion of liquid aerosols inside alveoli in the absence of ventilation, we paused ventilation at peak inspiratory volume, held the volume constant for two seconds, and observed the trajectories of droplets (projected onto the X-Y plane, with gravity acting in the Z-direction) that entered alveoli after ventilation was stopped (**Fig. 5f**). In the absence of ventilation, droplets traveled up to 6 μm and lingered in the observable airspace for typically 0.1–0.2 s (**Fig. 5g**), afterwards either impacting the alveolar wall or moving out of the objective focal plane. In a log-log plot of the mean squared displacement for droplets in air and helium, the value of the slope of a linear fit corresponds to the diffusivity of the particles in the airspace motion, where m > 1 corresponds to super-diffusive motion, m = 1 corresponds to diffusive motion, and m < 1 corresponds to sub-diffusive motion^63^ (**Fig. 5h**). Droplets traveling during paused ventilation exhibited diffusive motion (m ∼ 1) (**Fig. 5h**). Mean velocity (**Fig. 5i**), path length (**Fig. 5j**), and path persistence (total displacement divided by path contour length) (**Fig. 5k**) were also similar for all droplets regardless of gas composition.

Previously, aerosol deposition was found to be spatially heterogeneous from the lung apex to base and particles generally deposited in airways versus alveoli based on size but was generally assumed to be uniform between neighboring alveoli due to imaging limitations. Additionally, the distribution of liquid aerosols was poorly understood due to the confounding effect of rapid post-deposition spreading of liquid droplets disrupting the signal and preventing examination by histology. We applied the recently developed crystal ribcage coupled with ultrafast microscopy to challenge these assumptions and overcome limitations by directly visualizing aerosol distribution during real time, physiological breathing. We directly observed alveolar-scale spatial heterogeneity of aerosol deposition in the lungs, organized into a “mosaic pattern.” Aerosols (1-10 µm) deposited preferentially into “tile” regions while neighboring alveoli received negligible aerosols formed a “band” around the tile perimeter. Additionally, the mosaic pattern persisted in vivo and ex vivo regardless of ventilation scheme, aerosol material composition and molecular weight, lung age, and was conserved in murine and large animal models, with similar heterogeneity indicated in human lungs. These findings suggest that the differential distribution of inhaled pollutants, pathogens, and potentially gases within alveoli over the mammalian lifespan may drive heterogeneities in alveolar biology and immunology, shaping our fundamental understanding of lung function. While the biological and immunological implications of the mosaic pattern are beyond the scope of this study, they may influence the spatial heterogeneities associated with the development of lung pathologies linked to aging, such as respiratory infections, lung metastasis, emphysema, and fibrosis^64^. Additionally, while aerosols in the size range 1-10 µm are highly relevant for studies of disease pathogenesis and environmental pollution, the formation of the mosaic pattern for droplets outside of this range, such as gases, nano-scale aerosols, and large particulates, represent an important direction for future study.

In particular, aerosol redistribution at the lobar level has implications for both drug delivery and disease and injury models in the lung, as both material property and lung age determine whether only discrete groups of alveoli are exposed to inhaled materials, or whether the entire parenchyma is exposed, and over what timescale. Direct insight into the deposition and clearance (or retention) of these materials in the lung is crucial for understanding how lung immune response may be spatially and temporally organized given that only particular alveoli come into routine direct contact with harmful aerosols. Contemporary society is challenged by both the high prevalence of airborne microplastics, with increasing evidence of negative effects at all stages of biological organization, and threat of airborne pathogens. Understanding the transport physics of aerosols in the lung over time will inform public health strategies to improve environmental exposure controls and reduce harm, as well as revealing new therapeutic targets. Furthermore, in the context of lung development, future studies into the mosaic pattern could provide insight into gas transport heterogeneities as a potential driver of lung development, by utilizing emerging tools for optical imaging of gases such as oxygen partial pressure^65^. These insights may inform improved ventilation and therapeutic strategies to support normal lung development in premature infants^66^.

Lastly, we present the first direct visualization of single droplet transport during physiological breathing as aerosols enter the alveolar airspace. Previously, it was believed that aerosols on the scale of 5-10 µm would deposit in the upper airways and not travel to the level of alveoli. In consequence, only smaller particles would reach distal alveoli, where transport is dominated by a combination of diffusion and gravitational sedimentation dependent on particle size. However, we found that single aerosols as large as 10 µm do appear in the most distal alveoli in murine lungs. Moreover, if alveolar transport was diffusion dominated, droplets should land uniformly on all parts of the alveolar wall. In contrast, we found that during active ventilation, particles deposit preferentially on the surface of alveoli aligned with rigid body motion of the lung, indicating that inertial effects contribute to particle transport even at the alveolar level. However, in the absence of ventilation, aerosols travel with diffusive trajectories. These discoveries offer a unique opportunity to investigate the physics of aerosol transport beyond computational modeling, and to probe the interactions between aerosols containing pathogens, pollution, or therapeutics and the alveolar epithelium and resident immune cells.

## Acknowledgements

We thank the Boston University Neurophotonics Center, Boston University Micro Nano Imaging Core Facility, and Photonics Center for their generous support and access to their imaging resources, particularly the Nikon W1 SoRa spinning disk microscope acquired through NSF MRI Award #2215990. We thank New England Donor Services (Waltham, MA) for their assistance in human lung donor organ procurement and gratefully acknowledge the donors and their families for their selfless contribution.

## Author contributions

G.N.G. and H.T.N. conceived the project and wrote the manuscript; G.N.G. conducted the lung extraction and imaging experiments; H.A.K. performed and designed data analysis pipelines; V.T. optimized the PCLS protocol and performed data analysis; R.L. performed image processing and performed mathematical modeling of diffusion; R.B. developed the crystal ribcage and custom nebulizers; W.W. fabricated the mouse crystal ribcage; A.A.R. generated fibrosis mouse models; L.S. conducted lung extraction surgery for neonatal mice; S.B. and H.S. fabricated and characterized nanoparticles; A.B. designed and fabricated the crystal ribcage for neonatal mice; B.K. co-conducted human lung experiments; A.T. performed image processing; A.K. co-developed image processing and tracking pipelines. L.C. prepared tissue for histology; K.R. co-developed data analysis pipeline; F.D. synthetized and characterized nanobodies; M.V. prepared E. coli-GFP stock and growth characterization; G.N.G., R.B., L.S., L.C., K.R. maintained the mouse colony; C. D. conducted laser diffraction particle imaging studies; W.L. and M.H. designed and synthesized synthetic lung surfactant powder; G.L., M.R., L.H., J.P.M., W.L., M.H., W.M.S., J.P.B, and B.S. contributed to discussion on crucial aspects of the project; H.T.N. supervised the project.

## Methods

### Animal use ethics

All experimental procedures adhered to the ethical guidelines approved by the Boston University Institutional Animal Care and Use Committee. Mice were bred and housed at the Boston University Animal Science Center in a pathogen-free environment with ambient temperature, humidity, 12-hour light-dark cycles and unrestricted access to food and water. We did not make any exceptions to animal housing or handling protocols for this study.

### Mice

We used mice aged 24 hrs post birth (postnatal day 0) to 28 months of age, both male and female, for this study. To establish a colony of reporter mice for imaging lung structure, we purchased a breeding pair of transgenic B6.129(Cg)-Gt(ROSA)26Sortm4(ACTB-tdTomato,-EGFP)Luo/J (JAX #007676, Jackson Laboratory, Bar Harbor, ME), referred to as mTmG, to serve as the primary source of animals for imaging experiments. Male and female C57BL/6J were purchased from Jackson Labs when reporter fluorescence was not needed (JAX, #000664, Jackson Laboratory, Bar Harbor, ME). For aging studies, male and female C57BL/6 mice were acquired from the National Institutes of Health National Institute of Aging.

### Elastase-induced emphysema model

We generated a mouse model of emphysema in mTmG reporter mice by orotracheal administration of porcine pancreatic elastase (PPE) (EC143, Elastin Product Company). A single dose was administered according to a published protoco^48,49^ where 6 IU of PPE was dissolved in 100 μL of saline and delivered to the back of the tongue of anesthetized, vertically suspended mice. Mice were euthanized 3 weeks post injury for ex vivo lung imaging using the crystal ribcage. We prepared fixed samples of healthy and elastase-treated mouse lungs for cryotome slicing and hematoxylin and eosin (H&E) imaging of OCT-mounted slices and imaging by iHisto Inc. (Salem, MA).

### Melanoma metastasis

We utilized B16F10-luc-GFP mouse melanoma cells (Dana Farber Cancer Institute, Boston MA) to model melanoma metastasis to the lungs in C57BL/6J mTmG reporter mice or unlabeled C57BL/6J mice. We cultured the cells in DMEM (Corning) supplemented with 10% fetal bovine serum (Gibco) and 1% penicillin/streptomycin (Gibco). At approximately 80% confluency, we collected the cells, counted, and resuspended in complete medium prior to tail vein injection into mice at a concentration of 1 million cells in 200 µl of volume. We confirmed all cell lines to be mycoplasma-free (Mycoalert Plus Mycoplasma Detection kit, Lonza) before use in animals. We euthanized mice injected with cancer cells 1 to 10 days post-injection for microscopy. The largest tumor nodules examined were 2 mm in diameter, and we euthanized mice exhibiting labored breathing, a hunched posture, or ruffled fur due to tumor progression, as per BU IACUC guidelines.

### Methacholine airway constriction model

To probe the effect of airway constriction on mosaic pattern formation after aerosol inhalation, we delivered a dose of fluorescent liquid aerosols to the ex vivo mouse lung of reporter mTmG mice before and after delivering an intravascular bolus of methacholine to the ex vivo lung^47^. Prior to airway constriction, we nebulized 200 µL of DiD solution (1 mg/ml, in saline, **Table 1**) into the mouse lung by pressure-controlled positive pressure ventilation (120 breaths/min, end expiratory trachea pressure = 2 cmH_2_O, peak inspiratory trachea pressure = 18 cmH_2_O, tidal volume = 0.2 mL) over 2 minutes. Then, we imaged the baseline mosaic pattern by spinning disk confocal microscopy. Next, we injected a bolus of methacholine (200 µL, 10 mg/mL in saline) into the lung vasculature in combination with a bolus of cascade blue dextran (50 µL, 10 mg/ml) to act as a fluorescent tracer for methacholine distribution within pulmonary capillaries, as previously described^36^. We turned off perfusion flow after fluorescence was observed in pulmonary capillaries and allowed methacholine to incubate in the ex vivo lung for 10 minutes to cause airway constriction, confirmed by a decrease in tidal volume to less than 0.1 mL for the same pressure-control ventilation parameters. Then, we delivered a second dose of aerosols using 200 uL of nebulized fluorescein isothiocyanate (FITC) conjugated albumin (25 mg/mL in saline, **Table 1**) and imaged the resulting mosaic pattern after airway constriction, experimental design schematic in **Supplemental Fig. 9a**.

**Table 1.** Size description and source information for aerosolized materials.

| **Aerosolized material** | **Working concentration** | **Source** | **Molecular weight or characteristic size** |
| --- | --- | --- | --- |
| **Liquid formulations** |  |  |  |
| **Cascade blue hydrazide trisodium salt** | 10 mg/mL | C687, Thermofisher Scientific | 479 Da |
| **Lucifer Yellow CH, lithium salt** | 25 mg/mL in saline | 80015, Biotium | 457 kDa |
| **Evans blue (unbound dye)** | 10 mg/mL in saline | AAA1677418, Fisher Scientific | 961 Da (unbound), 70 kDa bound to albumin |
| **Cascade blue dextran** | 10 mg/mL | D1976, Thermofisher Scientific | 10 kDa |
| **Alexa Fluor 647 dextran** | 5 mg/mL | D22914, Molecular Probes | 10 kDa |
| **Bovine serum albumin-fluorescein isothiocyanate conjugate** | 25 mg/mL in saline | A9771, Sigma Aldrich | 70 kDa |
| **Fluorescein isothiocyanate-dextran** | 10 mg/mL in saline | 46945, Sigma Aldrich | 70 kDa |
| **Texas Red dextran** | 5 mg/mL in saline | D3329, Fisher Scientific | 3k Da |
| **Bovine serum albumin, Texas Red conjugate** | 5 mg/mL in saline | A23017, Molecular Probes | 70 kDa |
| **DiD (DiIc18(5); 1,1'-dioctadecyl-3,3,3',3'-tetramethylindodicarbocyanine, 4-chlorobenzenesulfonate salt), solid perchlorate** | 1 mg/mL in saline | 22034, ATT Bioquest | 960 Da |
| **DiO (DiOC<sub>18</sub>(3); Benzoxazolium, 3-octadecyl-2-[3-(3-octadecyl-2(3H)-benzoxazolylidene)-1-propenyl]-, perchlorate)** | 1 mg/mL in saline | 60011, Biotium | 882 Da |
| **DiD conjugated PLA-HPG nanoparticles** | 22.5 mg/mL | Provided by co-authors at Yale University | 100-200 µm |
| **Anti-fibronectin nanobody** | 2 mg/mL | Provided by co-authors at Boston University | 14-18 kDa |
| **Escherichia coli-GFP (E. coli-GFP)** | 10<sup>6</sup> CFU/mL | Strain 25922GFP (ATCC) | 1-2 µm in diameter |
| **Dry formulations** |  |  |  |
| **Fluorescent amino formaldehyde polymer dry powder, 1-5 µm microspheres** | Density 1.3 g/mL, ~50 mg per dose | Cospheric, FMG-1.3 1-5µm - 500mg | 1-5 µm |
| **Synthetic lung surfactant dry powder aerosol formulation** | ~20 mg per dose | Provided by co-authors at Virginia Commonwealth University | 1.2-1.3 µm |

### In vivo bleomycin-induced lung fibrosis

Male and female mTmG mice (2 months old) received 1.25 U/kg of Bleomycin in 50 µl saline via oropharyngeal administration as previously described^67^. Briefly, mice were anesthetized with isoflurane and when light plane anesthesia was achieved, the animals were placed on a vertical mouse intubation platform. Mouse tongue was gently pulled and held with forceps, bleomycin was administered into the distal part of the oral cavity (oro-pharynx) using a pipette, and the nose was gently closed until the liquid disappeared down the respiratory tract. The tongue was released back when the gasping of the animal completely stopped. Respiration was critically monitored, and the animals were placed in a horizontal position for recovery. Animals were returned to the cage after recovery from anesthesia.

### Crystal ribcage fabrication and development

The crystal ribcage is a transparent biocompatible, age- and strain-matched physiological environment for the ex vivo lung^36^. Detailed fabrication of the ribcage has been previously described^36^ and is summarized here. We used previously recorded microcomputed tomography scans of the mouse chest (C57B/6 scans were obtained courtesy of the Hoffman group at the University of Iowa^68,69,70^, and to define a 3D model of the chest cavity that we printed using a stereolithography printer (Form3, FormLabs, Massachusetts, USA). We polished the printed model to remove printing artifacts and then used it as a positive mold to thermoform the crystal ribcage. The base model was made age-specific by scaling the geometry by mouse lung volume variations with age^71^, for both neonatal and adult mice. The internal surface of the crystal ribcage was dip coated in iSurGlide Plus hydrogel (iSurTech, Minnesota, USA) following the manufacturer’s instructions to obtain a permanent, lubricious, and optically transparent hydrophilic coating on the crystal ribcage and provide a low-friction surface for lung ventilation. Briefly, we masked the outer surface of the ribcage with a two-part silicone material Ecoflex 00-30, Smooth-On, Pennsylvania USA), and we cleaned the masked ribcage with isopropanol and Kimtech wipes (Kimberly-Clark Professional, Georgia USA). Then, we dipped the ribcage into the iSurGlide solution for 30s before removing at a controlled rate of 3 mm/s and allowed the coated ribcages to degas and dry in a chemical fume hood for 5 minutes. We removed the silicone mask before the coated ribcage was placed on a rotary stage 3 inches away from an 8W 302nm UV-B light source to cure for 15 minutes. The final coated ribcages were stored in PBS to maintain hydration of the hydrogel coating and maintain tonicity while not in use.

### Ex vivo mouse lung preparation

Ex vivo lungs were prepared as previously described^36^. Briefly, we excised mouse lungs and ventilated them using the Kent Physiosuite Mouse Ventilator (Kent Scientific) or custom-built negative pressure ventilator via a tracheal cannula^37^. For experiments requiring lung perfusion, we perfused complete cell culture medium (RPMI + 10% FBS + 1% penicillin streptomycin) through the ex vivo lung, with the addition of fluorescent labels to visualize vascular lumens and circulating cells through two cannulas inserted into the pulmonary artery (PA) and left atrium (LA). Once the heart was cannulated, we loaded the whole heart-lung bloc into the crystal ribcage for microscopic imaging performed under controlled ventilation and perfusion. We regulated perfusion pressure by adjusting a 60-ml reservoir of the perfusate medium (serum-free RPMI, Gibco) to 5-15 cmH_2_O above the lung.

### Aerosol delivery to ex vivo and in vivo mouse lungs

Ex vivo lungs housed inside the crystal ribcage received liquid droplet aerosols that we nebulized using the Aerogen® Solo commercial nebulizer (Aerogen; Galway, Ireland) or Gülife ultrasonic mesh nebulizer (Amazon product) into the mouse trachea via a custom-built 3D printed adaptor. We nebulized liquid formulations containing 200 µL of dissolved fluorescent small molecules, nanoparticles, or whole pathogens, or sprayed amounts of dry powder (**Table 1**) into the trachea for 2 minutes during active ex vivo lung ventilation using a Kent Physiosuite Mouse Ventilator (Kent Scientific) (PPV) or by a custom-built negative pressure ventilator (NPV)^37^. Dye concentrations and type of nebulizer used were maintained consistently between biological replicates, see **Table 1**. Unless otherwise indicated in the figure or figure caption, lungs were ventilated at 120 breaths/min under volume-controlled ventilation with tidal volume = 0.25 mL, achieving a pleural pressure range of -2 to -10 cmH_2_O during negative-pressure ventilation, or +2 to +10 cmH_2_O tracheal pressure during positive pressure ventilation for 2 continuous minutes at 37°C environmental temperature. Negative inspiratory: expiratory ratio was 1:1, and 1:2 for positive pressure experiments using a consistent ventilation curve profile. For aerosol post-deposition redistribution experiments, ventilation was paused for 5 seconds every 5 seconds for the first 30 seconds to capture a high-resolution image of the whole lung lobe during active delivery, then held static at peak inspiratory volume for the remainder of the imaging time. During these experiments, the nebulizer was continuously delivering aerosols for the first 30 seconds before aerosol generation was stopped. To measure single droplet trajectories, images of single droplets traveling in the alveolar airspace were captured at 200 FPS during a 2-second pause every 4 breath cycles under air or helium ventilation.

For aerosolization in vivo, sedated mice were positioned supine on a flat cork surface, and a small incision was made in the trachea such that the mouse was spontaneously breathing through the trachea instead of the nasal passages. The mouse was then enclosed in a sealed plastic box. The nebulizer was attached to one wall of the box such that aerosols would be dispersed inside. While anesthetized, mice inhaled aerosols generated from the nebulizer by spontaneous breathing in intervals of 30 seconds with the nebulizer turned on, then 30 seconds off, for a total of 5 minutes. Afterwards, anesthetized mice were immediately sacrificed and lungs extracted for imaging inside the crystal ribcage.

### Aerosol delivery to ex vivo porcine lungs

We stored whole ex vivo porcine lungs (Research 87 Inc., Boylston MA) on ice for 1-2 hours prior to delivery for aerosol experiments. We recruited the left lung using a positive pressure clinical ventilator by increasing tracheal pressure from 8 to 20 cmH_2_O (over approximately 30 minutes) through a custom-built 3D tracheal cannula with nebulizer attachment, suspended in the air inside a 37°C heated chamber with a commercial humidifier to maintain tissue hydration, without using a crystal ribcage. We nebulized 5 mL of Evans blue (1 mg/ml) or DiD dye (1 mg/ml) in saline into the pig lungs during positive pressure ventilation (30 breaths/min, positive end expiratory pressure (PEEP) = 2 cmH_2_O, peak inspiratory pressure (PIP) = 20 cmH_2_O) over approximately 30 minutes. After aerosol delivery, we held the lungs inflated at 15 cmH_2_O and imaged using an upright Nikon stereomicroscope with 0.67, 1x, and 2x objectives, and Nikon X1 spinning disk confocal microscope with 1x, 2x, and 4x objectives.

### Aerosol delivery to ex vivo human transplant-rejected lungs

We received whole ex vivo human lungs not suitable for transplant 6-12 hours after operation from New England Donor Services (Waltham, MA). Patient 1: male, 71 years old, history of smoking and COPD. Patient 2: female, 46 years old, history of depression and stroke. During operation, the surgical team perfused lungs with saline and stored the deflated tissue in a saline bath on ice for transport. For ex vivo aerosol delivery, we recruited the left lung using a positive pressure ventilator by increasing trachea pressure from 8 to 20 cmH_2_O (over approximately 30 minutes) through a custom-built 3D tracheal cannula with nebulizer attachment. We suspended the whole organ in the air inside a 37°C heated chamber with a commercial humidifier to maintain tissue hydration, without using a crystal ribcage. We nebulized 5-10 mL of DiD dye (1 mg/ml) in saline into the lungs during positive pressure ventilation (30 breaths/min, positive end expiratory pressure (PEEP) = 2 cmH_2_O, peak inspiratory pressure (PIP) = 30 cmH_2_O) over approximately 30 minutes. We used aerosol delivery to model the intrapulmonary delivery of immunosuppressive therapeutics to the human lung. After aerosol delivery, the lung pleural surface was imaged using a Nikon X1 spinning disk confocal microscope with 1x, 2x, and 4x objectives. This project was determined to be IRB-exempt by the Charles River Campus Institutional Review Board at Boston University as the procedures and donor organs do not meet the definition of “human subjects”, defined in the U.S. Department of Health and Human Services, Office for Human Research Protections regulation 45 CFR 46.102(l).

### Nanoparticle fabrication

We synthetized PLA-HPG nanoparticles (**Table 1**) as previously described^39^. DiD loaded PLA-HPG nanoparticles were fabricated as we previously described ^39,72,73^ with minor modifications. Briefly, we dissolved 45 mg of PLA-HPG in 2.4 mL ethyl acetate (EA). We dissolved DiD in 600 µL of DMSO and added it to PLA-HPG in EA to reach a DiD to PLA-HPG mass ratio of 0.2%. We added the polymer and dye solution to 4 mL of deionized (DI) water under vortex. We probe sonicated the emulsion for three cycles, each lasting 10 seconds, with a brief rest on ice between each sonication. We diluted the final emulsion to 30 mL with DI water and placed it under rotary evaporation for 30 min to remove EA. We transferred the nanoparticle solution to Amicon Ultra Centrifuge Filtration Units (100kDa molecular weight cutoff) and washed with cold DI water for two cycles to remove DMSO. We resuspended the resulting particles to 2 mL in a 7.5% sucrose solution and froze to -20°C. We used a small amount (10 µL) of the particle solution to evaluate the formulation by dynamic light scattering (DLS) using a Malvern Zetasizer Nano: particle size = 108.9 ± 1.8 nm, polydispersity index = 0.23 ± 0.03, Zeta potential = -10.7 ± 0.06 mV.

### Fluorescent nanobody synthesis

We cloned and expressed anti-fibronectin nanobody as previously reported^38^. To label the nanobody, we added tris(2-carboxyethyl)phosphine hydrochloride (TCEP, final concentration 5 mM) to 1 mg nanobody solution and incubated at room temperature for 30 min. We dissolved IVISense 645 MAL Fluorescent Dye (VivoTag, Revvity) in DMSO to 10 mg/mL and added it to the nanobody at a 10:1 molar ratio of dye to nanobody. We incubated the resulting solution at room temperature in dark for 4 h then purified the product by ultracentrifugation (MilliporeSigma™ Amicon™ Ultra-0.5, MWCO = 3 kDa) at 12,000 x g for 10 min and washed with cold PBS 3 times. We determined the final concentration of the fluorescent nanobody by measuring its UV absorbance at 280 nm.

### Synthetic lung surfactant dry powder formulation and dry particle aerosol delivery

We prepared the synthetic lung surfactant dry powder aerosol formulation (**Table 1**) fluorescently labeled with Oregon Green using similar methods to those previously described^41^. Briefly, we spray dried feed dispersions containing DPPC, POPG, mannitol, sodium chloride, L-leucine combined at a ratio of 55:7.5:9:6:20% w/w using the Buchi Mini Spray Dryer S-300 (Büchi Labortechnik AG, Flawil, Switzerland). We added 0.5 % of Oregon Green DHPE, (O12650, ThermoFisher Scientific, Waltham, MA) (0.5% of the total solid content) for fluorescent microscopy of dry powder aerosol. We spray dried the formulation in the open mode configuration using an inlet temperature of 80°C, formulation feed rate of 2 ml/min and drying gas flow of 30 m^3^/hr. We collected spray-dried powders into glass vials and stored them in a desiccator (<20% RH) at 5°C. For delivery of surfactant aerosols to the ex vivo mouse lung, 1-3 doses of 10-15 mg of powder each were loaded into a Small-Animal Air-Jet Dry Powder Insufflator^41^ ^40^ and delivered as a bolus aerosol to the ex vivo lung during negative-pressure ventilation, ventilating the lung for 2 min between doses.

For delivery of dry microplastics to the mouse lung, 10 g of dry microplastic beads (Table 1) were loaded into a glass Hamilton syringe (3 mL) with the syringe tip fitted with an 8 μm plastic mesh cut from a Thermo Scientific™ Nunc™ Polycarbonate Membrane Insert (Thermofisher Scientific). The mesh served as a filter and was connected to the lung trachea via a 1/8” ID silicone by a T-connection, with the other end connected to a 4L sealed air reservoir. While the lung (housed in the crystal ribcage) was negative pressure ventilated, the syringe was depressed 1.5 mL volume once every second to push the microplastic beads through the fine mesh and disperse aerosolized particles into the lung trachea. Aerosol administration lasted for 2 minutes.

### Laser diffraction particle size quantification

Nebulizer droplet size characterization performed by laser diffraction was done using a Malvern Spraytec® (Malvern Instruments, Ltd., Worcestershire, UK) with Spraytec® 4.00 software. The nebulizer exits were placed 1.5 cm from the laser beam and 6 cm from the detector in an open bench configuration. Nebulizers were loaded with 75 uL samples to generate an aerosol for a 10 second sampling period. A 300 mm lens with a laser-to-detector distance of ∼23 cm was employed with a 1 kHz acquisition rate. Automatic alignment and background subtraction were performed for each analysis. Data processing was concentration weighted.

### Preparation and characterization of Escherichia coli-GFP (E. coli-GFP)

We purchased *Escherichia coli*-GFP (E. coli-GFP) from ATCC (25922GFP). We cultured bacteria overnight at 37°C, 5% CO_2_ in 10 ml of THY broth. The next morning, we prepared 1:100 dilutions from the bacterial culture and incubated the cultures at 37°C, 5% CO_2_ for 8 hours. We checked bacterial cultures for growth and prepared aliquots in UV crosslinked Eppendorf tubes at a final concentration of 35% glycerol. We grew bacterial preps from aliquoted stock overnight at 37°C, 5% CO_2_ on blood agar plates. We selected single colonies and brought them to an optical density of 0.300 at 600nm in sterile saline. We prepared dilutions of 1:10, 1:15, 1:20 and 1:25 and plated 100 μl of each on blood agar and grown overnight at 37°C, 5% CO_2_. We then counted single colonies the next morning to determine CFU concentrations. A 1:10 dilution resulted in the desired final concentration of ∼ 1×10^6^ CFU and would be used for downstream aerosol experiments.

### Whole lung microscopy

We imaged ex vivo lungs within the crystal ribcage while maintaining environmental temperature at 37°C during quasi-static inflation or dynamic ventilation using an upright Nikon fluorescent stereomicroscope equipped with a green florescence filter and UV excitation lamp (NightSEA), Nikon CSU X1 upright spinning disk confocal, and Nikon CSU-W1 SoRa inverted spinning disk confocal microscope (NIS Elements software) with 1x, 2x, 4x and 10x, 40x silicone immersion, and 60x water immersion objectives. Microscopy data was reviewed and analyzed using NIS Elements, FIJI, and MATLAB R2024a. Two-photon microscopy for imaging collagen fibers by second harmonic generation (SHG) was done using a Bruker two-photon upright microscopy equipped with a 16x water immersion objective using Prairie View acquisition software.

### Precision-cut lung slicing and imaging

We excised mouse lungs following the ex vivo lung preparation procedure, then administered aerosolized DiD (20 uL,10 mg/mL) over 2 minutes using negative-pressure ventilation. The dye was allowed to permanently label cellular membranes for 1 hour, and then we filled the lung airspaces with 0.6 ml of 2% low-melting temperature agarose (BP16525, Fisher Scientific) dissolved in phosphate buffer saline (PBS) mixture and warmed to 37-40°C. After setting the agarose, we fixed the lung overnight submerged in 10% formalin, then triple-washed in PBS. We set the lung in agarose for sectioning using a Compresstome tissue slicer (VF-310-0Z, Precisionary Instruments, Ashland MA). We sliced the lungs from the pleural surface of the lung to toward the proximal core at a speed of 30% and oscillation of 60% of the maximum settings to obtain 400 µm thick slices. We collected the slices in a glass-bottom well plate for optimized imaging and set the slices in agarose to immobilize them during imaging. Next, slices were treated with thiodiglycol (TDE) for 4 hours following a steadily increasing concentration gradient (20%, 40%, 60%, 80%, in PBS, 1 hour per concentration) to match the index of refraction between the lung tissue and surrounding agarose, to maximize imaging depth. We used 1x, 4x, 10x and 20x objectives on the Nikon X1 dual camera spinning disk microscope to image the slices.

### Alveolar strain measurement

We measured tissue displacements using a symmetric, diffeomorphic implementation of deformable image registration provided by the ANTs registration library: 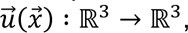 of the tissue at the cellular length scale at peak lung inflation and deflation (**Extended Data Fig. 9**). Using a cubic smoothing spline, we then evaluated the gradient derivative of the displacements, 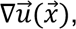 and recovered the linear approximation of the strain, 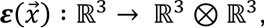 using the following equation.

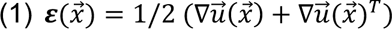

To select the frames of the image that correspond to inflation and deflation, we found the frames that exhibit the largest variance in intensity, which corresponds in practice to the frames that are in focus.

### Segmented alveolar area measurement

We acquired microscopy images using a 4x or 10x objective and took the maximum projection in the z direction across all pressure conditions in FIJI-ImageJ processing software. We identified alveoli from bands and tiles that stay within the field of view at all quasi-static steps or at both peak inhalation and peak exhalation points for dynamic ventilation data, before measurements were taken. Alveolar area was determined by using FIJI-ImageJ to calculate the area within annotations drawn by manually segmenting the alveolus-air interface for several alveoli (**Extended Data Fig. 9**). These average measurements were recorded for all alveoli at different pressures, then processed in MATLAB R2021b to generate plots and compute statistics across all groups.

### Trachea imaging window

We fabricated the trachea imaging chip from a silicon tube with a rectangular (∼1mm x ∼2mm) cutout, setting a plastic coverslip cut to the shape of this area to close the opening using Ecoflex™ FAST two component quick-setting silicone. Then, we secured the imaging chip between the nebulizer and trachea cannula and delivered aerosolized Lucifer Yellow dye dissolved in saline (25 mg/mL) to the lung under negative- or positive-pressure ventilation in the crystal ribcage. Using the trachea window, we imaged droplets as they entered the trachea using a Nikon W1 dual camera spinning-disk confocal microscope at 4x or 10x magnification, 5-10 ms frame interval. After imaging through the window, we moved the objective to the surface of the crystal ribcage to image the droplets arriving in alveoli during aerosol delivery in the same lung, with the same imaging settings.

### Single aerosol size quantification

We used FIJI (version 2.14.0) to export quantifiable frames during a two-second inspiratory hold from timeseries images of aerosol transport during inspiration. We developed a custom image processing pipeline to detect and analyze droplet edges using MATLAB (version R2023a Update 5). Our MATLAB script loaded TIFF stacks and normalized each frame to standardize intensity values. To account for variations across the image sequence, we employed histogram matching, ensuring consistent intensity distributions throughout the stack. A thresholding step was then applied to create a binary mask, isolating the brightest structures in the images. To mitigate noise while preserving edge information, we applied a 3D median filter to the matched image stack. We performed edge detection on each denoised image using the Sobel method with an empirically determined threshold. For object identification, we utilized MATLAB’s ‘regionprops’ function to extract bounding boxes for each potential object in the edge-detected images. An iterative algorithm was implemented to merge overlapping bounding boxes, resulting in a set of non-overlapping boxes that more accurately represented distinct objects. For visual inspection and validation, we created composite RGB images combining the original grayscale images with the edge-detected images and overlaid the final bounding boxes. This process was applied to each image in the stack individually, allowing for analysis of temporal changes in edge detection and object identification throughout the time series to detect droplet boundaries. Areas inside these boundaries were used to estimate droplet diameters.

### Single aerosol flux in variable tidal volume and variable respiratory rate

We used the trachea imaging window protocol above to image droplets passing from the nebulizer into the mouse trachea during active ventilation using a Nikon W1 dual camera spinning-disk confocal microscope at frame interval of 5 ms (200 FPS) at 4x magnification. For each tidal volume or respiratory rate condition, we exported peak inspiration frames from 10-12 ventilation cycles. On these frames, we applied manual counting using FIJI’s (version 2.16.0) cell counter tool to get total droplet count and divided this count by the number frames used, to report droplet number normalized by the number of breath cycles quantified for each tidal volume or respiratory rate condition.

### Single aerosol angle of impact quantification

We collected dynamic data of single aerosol transport during active ventilation using a Nikon W1 dual camera spinning-disk confocal microscope at frame interval of 5 ms (200 FPS) at 10x magnification. Trajectory angles of single droplets were measured from the center of the alveolus to the location of impact on the alveolar wall, measured using the line ROI tool in FIJI (version 2.14 1.54f). Droplets that did not impact alveoli within the imaging field of view (due to the constant respiratory motion) were not quantifiable.

### Single aerosol trajectory measurements

We collected dynamic aerosol inhalation microscopy data using the Nikon W1 dual camera spinning-disk confocal microscope. We collected time series data for aerosolized dye particles entering the alveolar space via nebulizer using a frame interval of 5 ms. To mitigate motion artifacts and ensure precise droplet tracking, we developed an image stabilization pipeline implemented in Python 3.9, utilizing NumPy (1.21.0), SimpleITK (2.1.0) ^74^, OpenCV (4.5.3), and tifffile (2021.8.30) libraries. We separated the acquired data into two TIFF stacks: a structure channel and an aerosol channel, which were read using the tifffile library.

We first converted each frame to float32 format and normalized. The frames then underwent preprocessing to enhance feature detection, including conversion to 8-bit format and application of Contrast Limited Adaptive Histogram Equalization (CLAHE)^75^ with a clip limit of 2.0 and tile grid size of 8x8. For feature detection, we employed the Oriented FAST and Rotated BRIEF (ORB) algorithms^76^, configured with 1000 features per frame, a scale factor of 1.2, and 8 levels. We used a brute-force matcher with Hamming distance and cross-checking for feature matching between consecutive frames.

We used the top 15% of matches, sorted by distance, to estimate an initial affine transform using OpenCV’s estimateAffinePartial2D function. In cases where this function failed, we used an identity transform instead. This initial transform guided a more refined B-spline registration using SimpleITK^77^. We initialized a B-spline transform with a mesh size of 10x10 for the 2D images, allowing for local deformations. For the registration method we utilized a correlation metric and the Limited-memory Broyden-Fletcher-Goldfarb-Shanno algorithm (L-BFGS-B) optimizer^78^ with gradient convergence tolerance of 1e-5, maximum 100 iterations, 5 corrections, 1000 function evaluations, and a cost function convergence factor of 1e7. We used linear interpolation for image resampling.

We implemented comprehensive error handling to ensure pipeline stability. For frames with insufficient features (<4 good matches), we skipped the registration and retained the original frame. In cases of registration failure, the algorithm defaulted to the initial affine transform. We saved the stabilized image sequences as 8-bit TIFF stacks using the tiff file library: stabilized structure channel, stabilized aerosol channel, and a composite visualization. We used the TrackMate plugin^79^ for FIJI (version 2.14.0)^80^ (Configuration: Hessian detector, Kalman tracker) to acquire trajectories of droplets arriving into the alveoli. We validated the trajectories by eye to filter out faulty tracks. We then exported 2-D Trajectories into a custom MATLAB (version R2023a Update 5) script to perform mean squared displacement (MSD) measurements. We were confined to tracking trajectories projected to the XY plane as the microscope cannot capture multiple z-planes in the time needed to capture single droplet motion. We tracked droplets until they impacted on the alveolar wall or moved out of the objective focal plane.

### Post-deposition aerosol transport quantification and diffusion coefficient estimation

We excised mouse lungs following the above lung preparation procedure, then nebulized a mixture of fluorescent dyes (see **Table 1**) into the lungs during ventilation for 15 breaths, followed by a 5 second hold at maximum inspiratory volume for image capture, repeated for 3-4 ventilation-pause cycles, before stopping aerosol delivery and maintaining lung ventilation paused at maximum inspiratory volume for an additional 10 min to several hours. We collected time-series data with 2x magnification using a Nikon SoRa W1 spinning disk confocal microscope. We loaded recorded time-lapses into FIJI (version 2.14.0) and drew rectangular ROIs across the diameter of the mosaic mesotiles with uniform short-side length. We exported fluorescence intensity across these ROIs at each time point in the form of distance across the long side of the ROI vs. mean intensity across the short side of the ROI. We loaded the profiles into a custom MATLAB (version R2023a Update 5) script to determine the full width at half maximum (FWHM) values of each mosaic tile across time after applying a min-subtraction correction to the data, which was necessary as mean brightness in the image steadily increased over time due to aerosol redistribution deeper inside the lung emerging to the surface. We averaged the FWHM values across tiles from the same mice and then reported average across all mice.

Upon measuring profiles of the image’s intensity, which we assume to be proportional to the concentration of solute, we observe that the intensity profile at each time step after administration of aerosol resembles an isotropic Gaussian distribution whose variance in the x-direction increases linearly with time. For Fick’s second law of diffusion in an unbounded domain, Green’s function precisely matches this observation. In particular, the concentration of solute, *φ*(*r*, *t*), evolves as a function of the diffusion coefficient *D*, radial distance *r* from the initial unit impulse and time *t*, as follows:

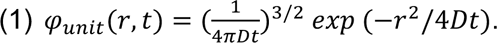

The radius, *r*_*FWHM*_, at which the distribution achieves half of its maximum value occurs where the concentration of solute, *φ*_*unit*(_(*r*, *t*), equals *φ*_*unit*(_(0, *t*)/2, which happens when the exponential term equals 1/2.

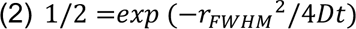

Solving for *r*_*FWHM*_^2^ yields the squared radius at which the concentration equals half its maximum value as a function of the diffusion coefficient and time.

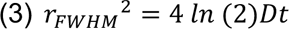

Given this equation, we determine the diffusion coefficient by finding the slope of *r*_*FWHM*_^2^ versus time.

### Statistics and reproducibility

We did not use statistical methods to predetermine experimental sample size. We did not exclude data from analysis. We did not blind investigators to allocation during experiments and did not randomize experiments. Data are presented as mean ± SEM. We calculated statistical values as a one-tailed Student’s *t*-test with Welch’s correction or one-sided ANOVA unless otherwise noted to determine significant *P* values between groups of data, considered significant for *P* < 0.05. *P* values are reported on figures to describe the statistical trends of the data. We generated plots and performed statistical analyses in MATLAB R2023a, R2024b, Microsoft Excel 16.91, and GraphPad Prism 10.3.1.

### Data availability

All data are available in the main text, Extended Data section, and Supplemental Videos section. Correspondence related to raw data and materials should be addressed to H.T.N.

