## Supplemental figures and videos for "Mosaic pattern: lung functional heterogeneity at the alveolus level"

### Supplemental videos

**Supplemental Video 1 |** Aerosol delivery to the lung inside crystal ribcage during real time microscopy

**Supplemental Video 2 |** Dynamic inhalation of aerosols produces mosaic pattern

**Supplemental Video 3** | Single droplets accumulate only in specific alveoli over time during active aerosol delivery, leading to the formation of the mosaic pattern by direct droplet deposition

**Supplemental Video 4** | Local perfusion through pulmonary capillaries does not spatially correlate with the mosaic pattern

**Supplemental Video 5 |** Post-inhalation redistribution of soluble dyes of variable molecular weight in young and aged mice

**Supplemental Video 6 |** Dynamic redistribution of a single aerosol across the alveolar septum

**Supplemental Video 7 |** Real-time imaging of single aerosols at the trachea and inside alveoli

**Supplemental Video 8 |** Arrival and deposition of aerosols on the alveolar wall

**Supplemental Video 9 |** Droplets deposit on the alveolar wall in directions aligned with lung rigid body motion during ventilation

### Extended Data Figures

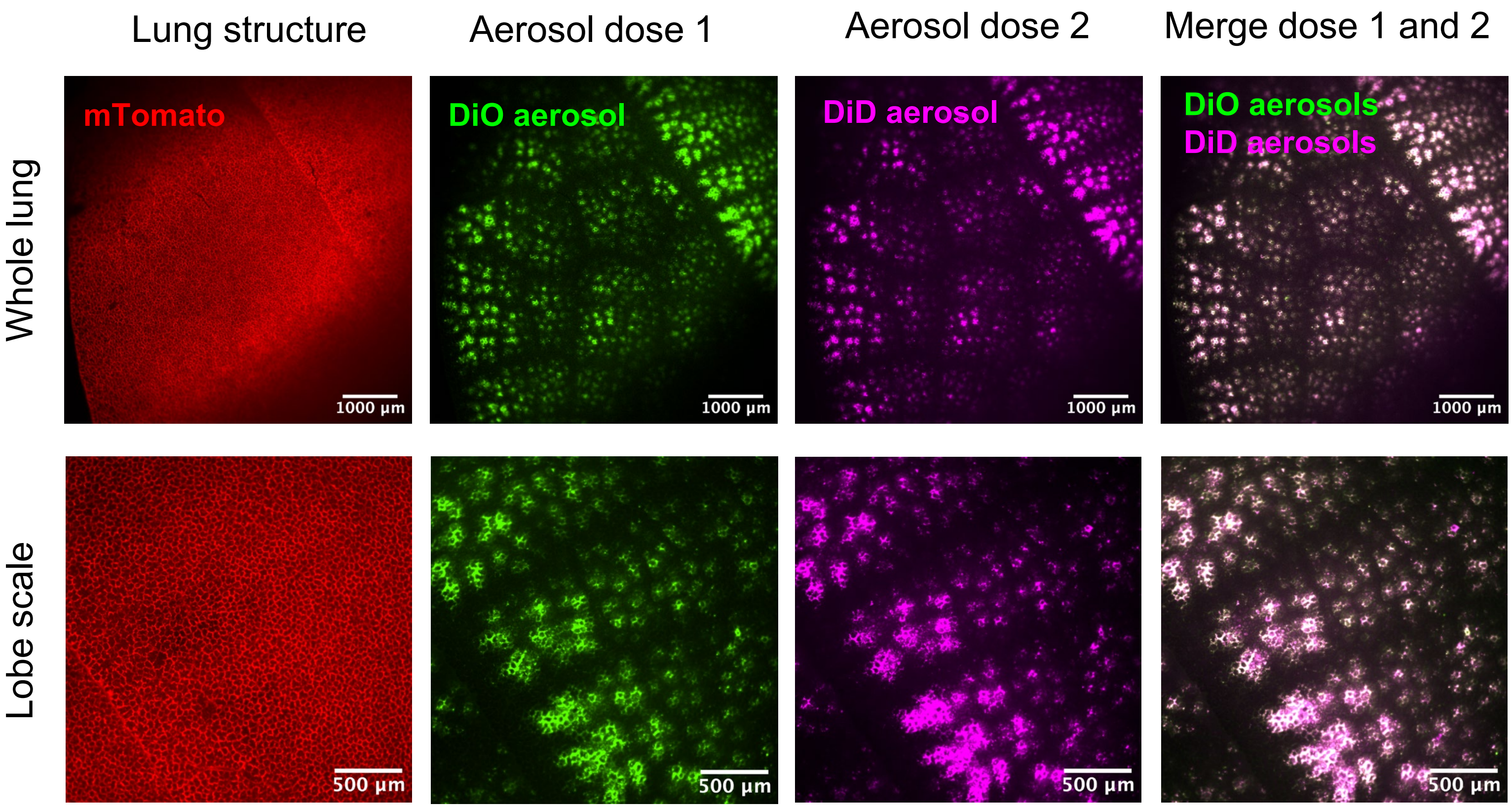


**Extended Data Figure 1 | The mosaic pattern is conserved across multiple ex vivo doses.** The same mosaic pattern is produced by liquid aerosols with different fluorescence delivered in sequence to the same mouse lung, where macrotile, mesotile, and microtile organization is preserved, n = 3 independent mouse lungs.

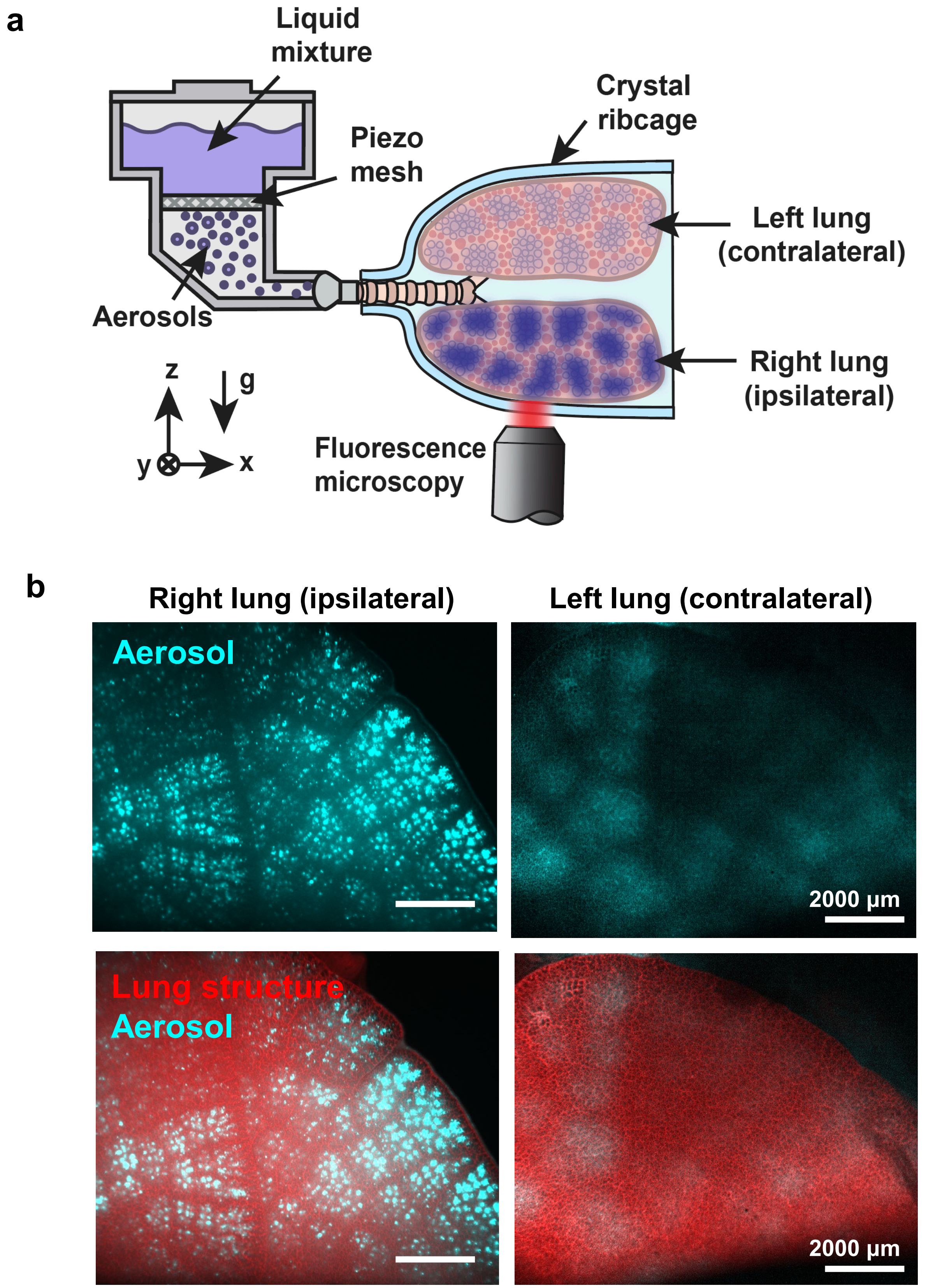


**Extended Data Figure 2 | Gravity dependence of the mosaic pattern**. (**a**) Aerosols deposit on the surface of the lung aligned with the direction of gravity after 2 min of aerosol inhalation. (**b**) The mosaic pattern is brightest on the right lung which is aligned with gravity during aerosol delivery (n = 3 independent mice).

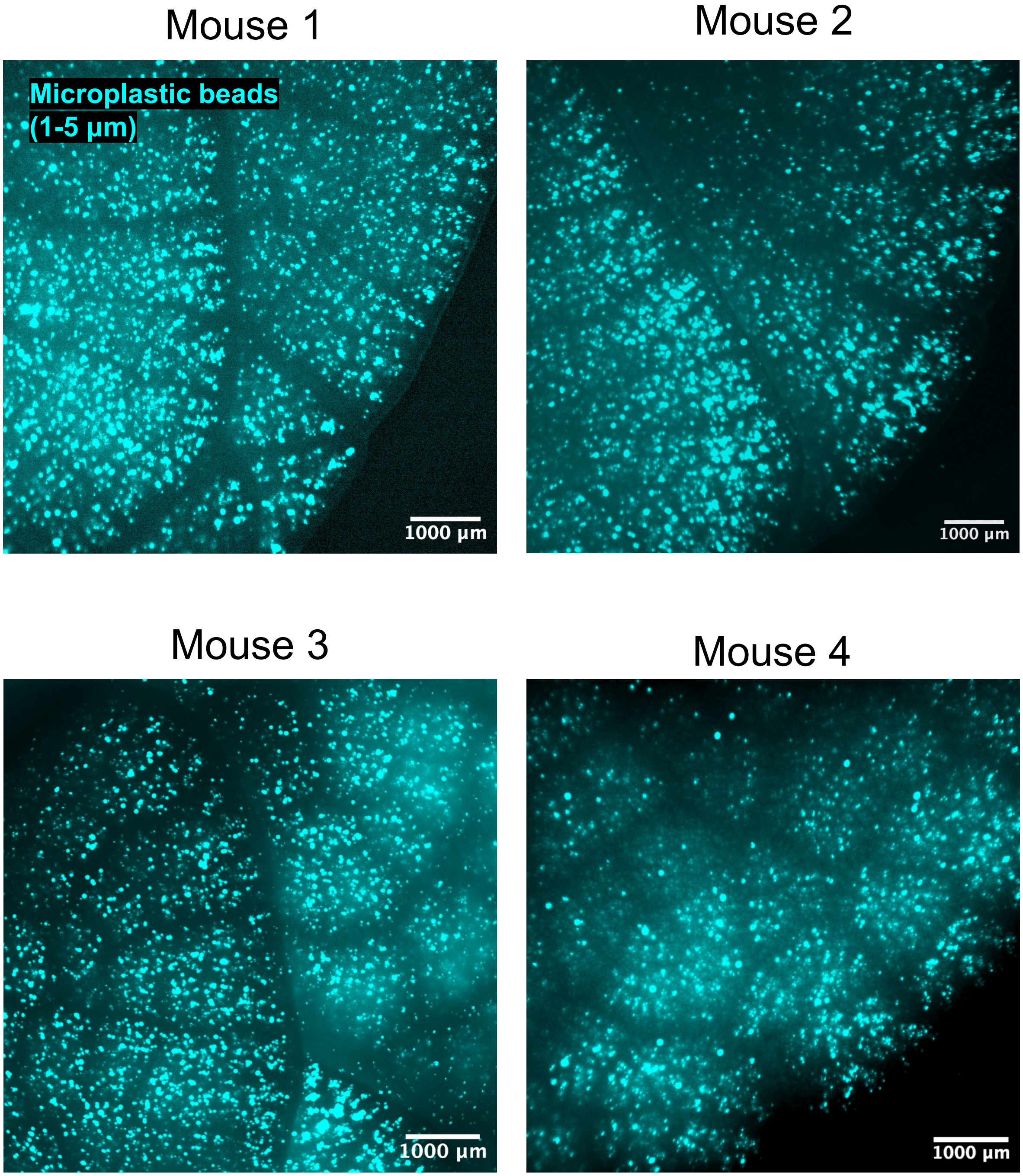


**Extended Data Figure 3 | The mosaic pattern is produced after inhalation of dry power aerosols**. Inhalation of dry fluorescent microplastic beads (1-5 μm) produces a mosaic pattern similar to the pattern produced by liquid droplet inhalation, n = 4 independent mouse experiments.

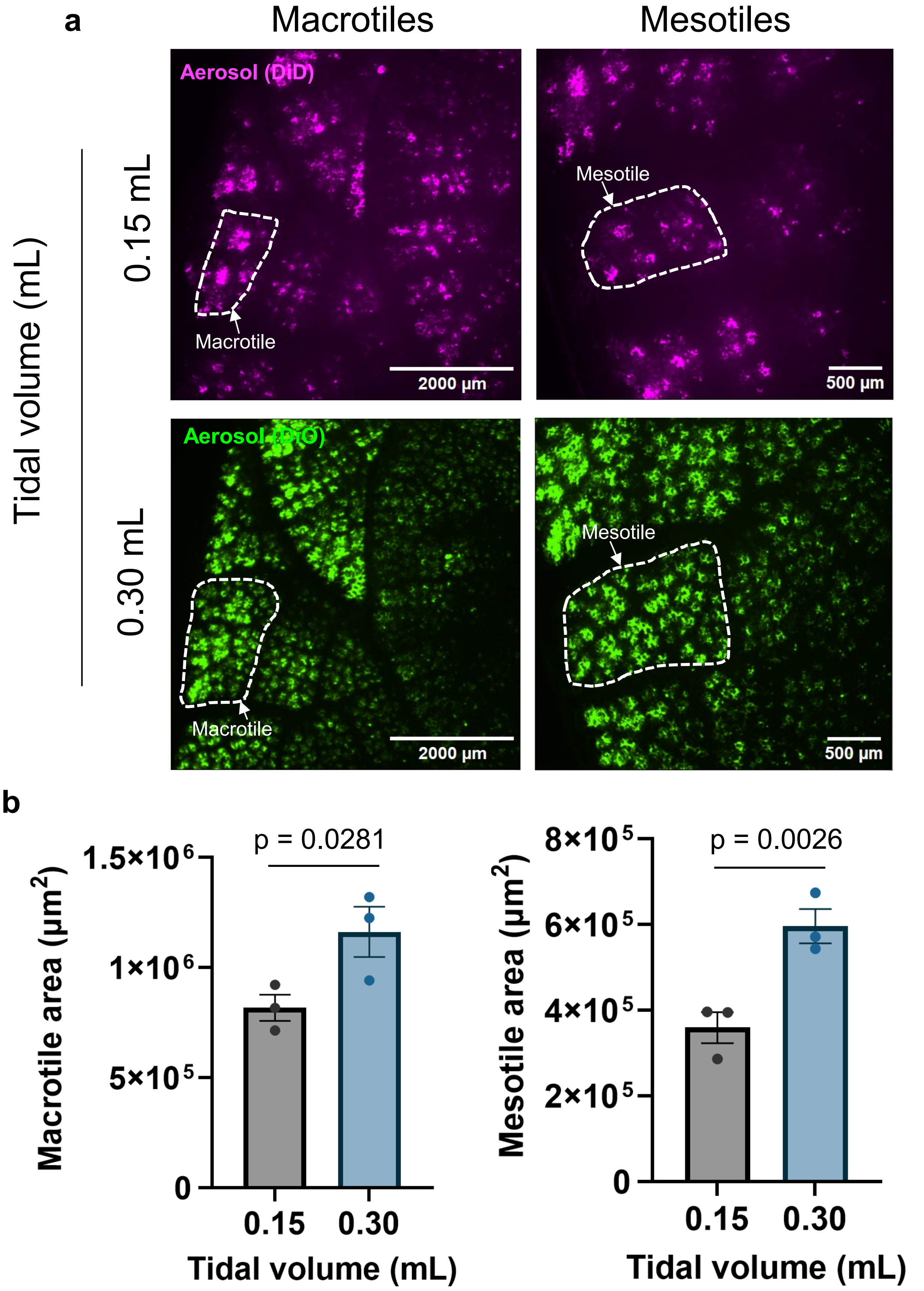


**Extended Data Figure 4 | Tidal volume causes an increase in mosaic pattern size across spatial scales**. (**a**) Representative images of mosaic pattern macro- and mesotiles formed after inhalation of aerosols under low tidal volume (0.15 mL) and high tidal volume (0.3 mL) conditions, keeping all other respiratory parameters consistent. (**b**) Mosaic tile area increases with increasing tidal volume at both the macro- and mesotile scale, N = 3 independent mouse experiments. Data are mean ± SEM, p-value calculated by paired t-test.

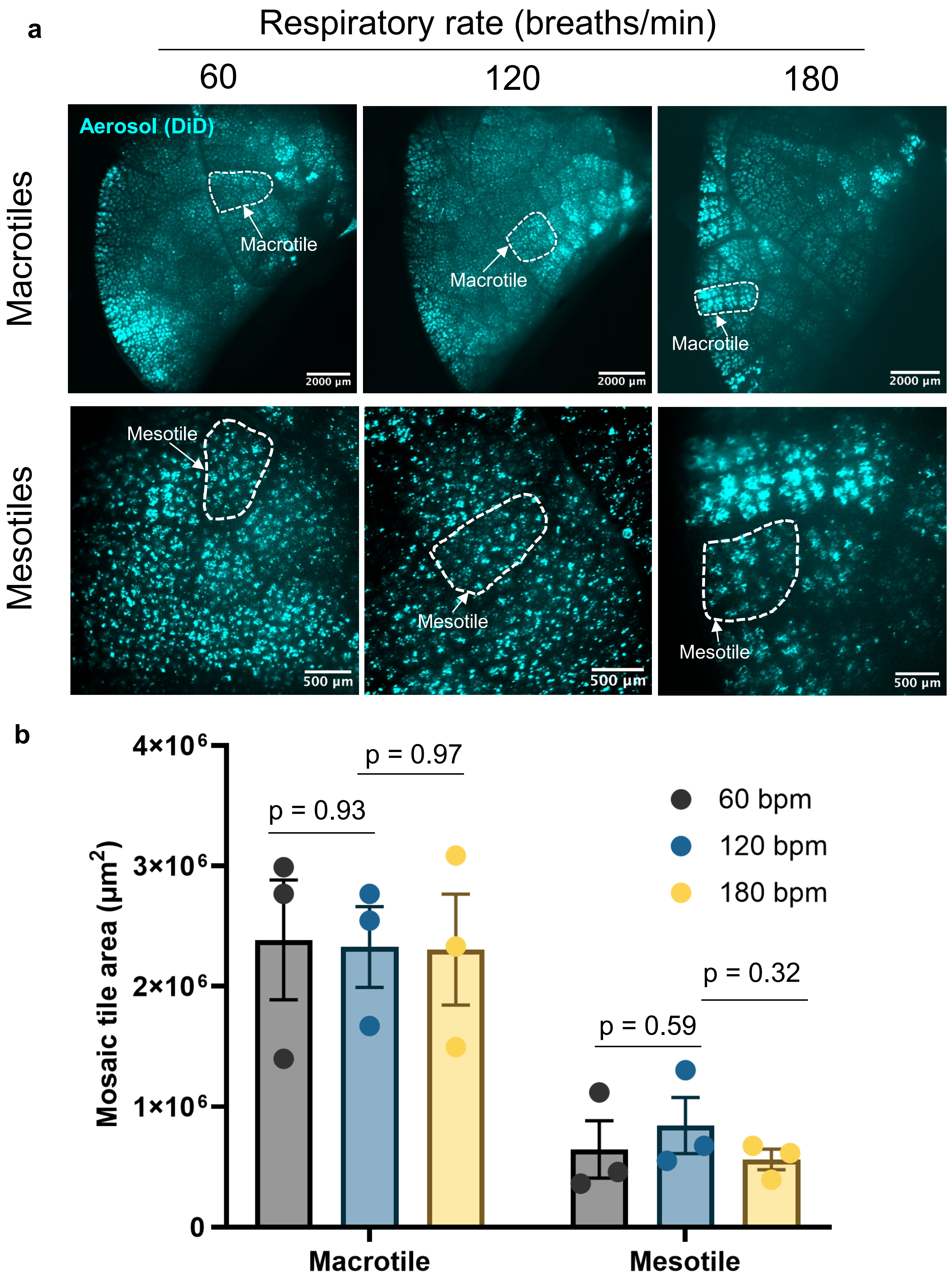


**Extended Data Figure 5 | Respiratory rate does not alter mosaic pattern tile size.** (**a**) Representative images of mosaic pattern macro- and mesotiles formed after inhalation of aerosols at increasing respiratory rates: 60, 120, and 180 breaths/min (bpm), holding all other respiratory parameters constant. (**b**) Mosaic tile area remains similar regardless of respiratory rate at both the macro- and mesotile scale, N = 3 independent mouse experiments. Data are mean ± SEM, p-value calculated by unpaired t-test.

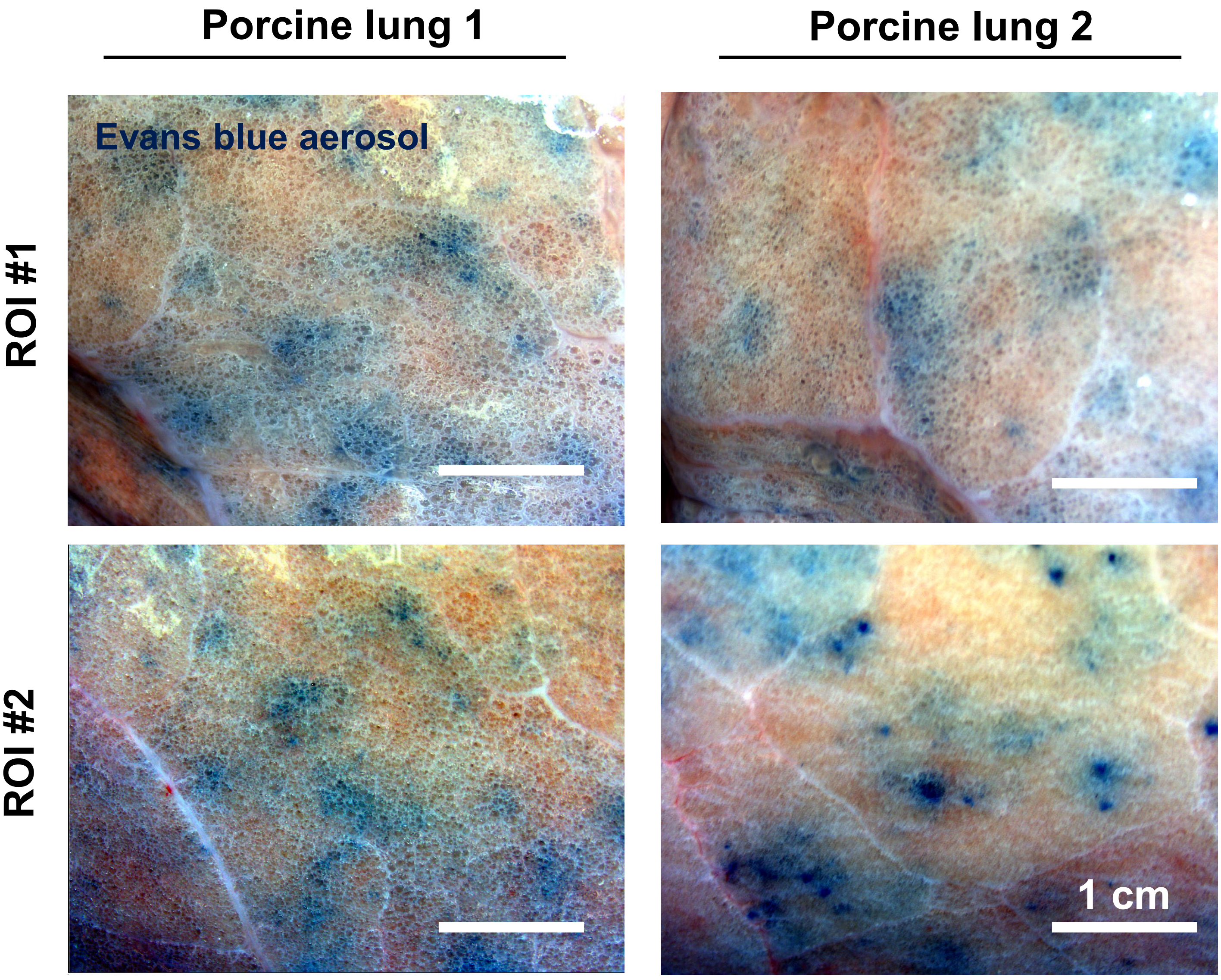


**Extended Data Figure 6 | Mosaic pattern heterogeneity is conserved in large animals**. Tile-like heterogeneity among neighboring populations of alveoli is visible in porcine lungs (n = 2 independent porcine lungs).

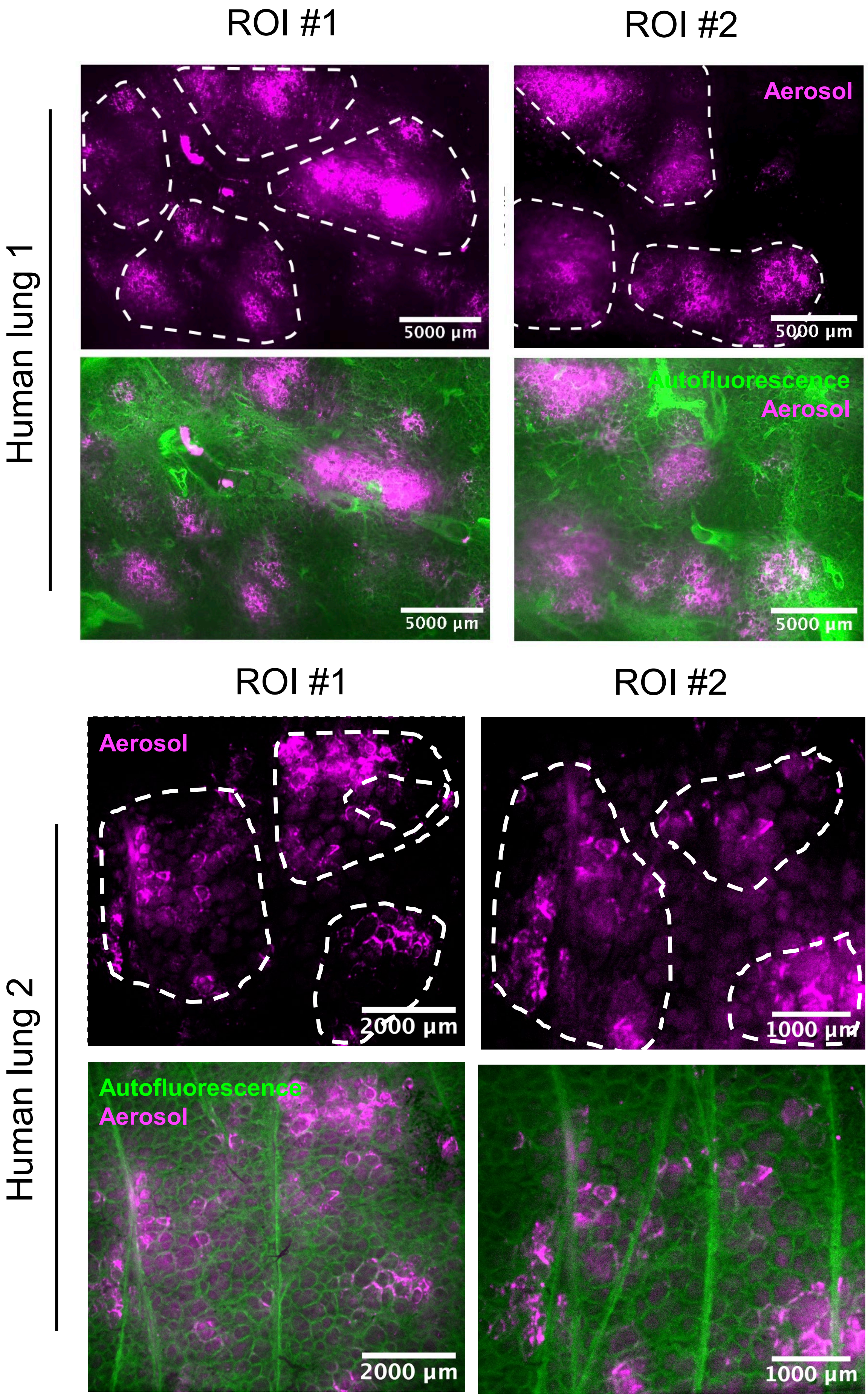


**Extended Data Figure 7 | Mosaic-like heterogeneity is present after aerosol delivery to human transplant-rejected lungs.** Large mosaic tile-like clusters of aerosol deposition (white dotted lines) are visible at the pleural surface using spinning disk confocal microscopy, indicating that intrapulmonary delivery of therapeutics (e.g. immune suppressants) to the human lung is heterogeneous at the alveolar level (n = 2 independent human lungs).

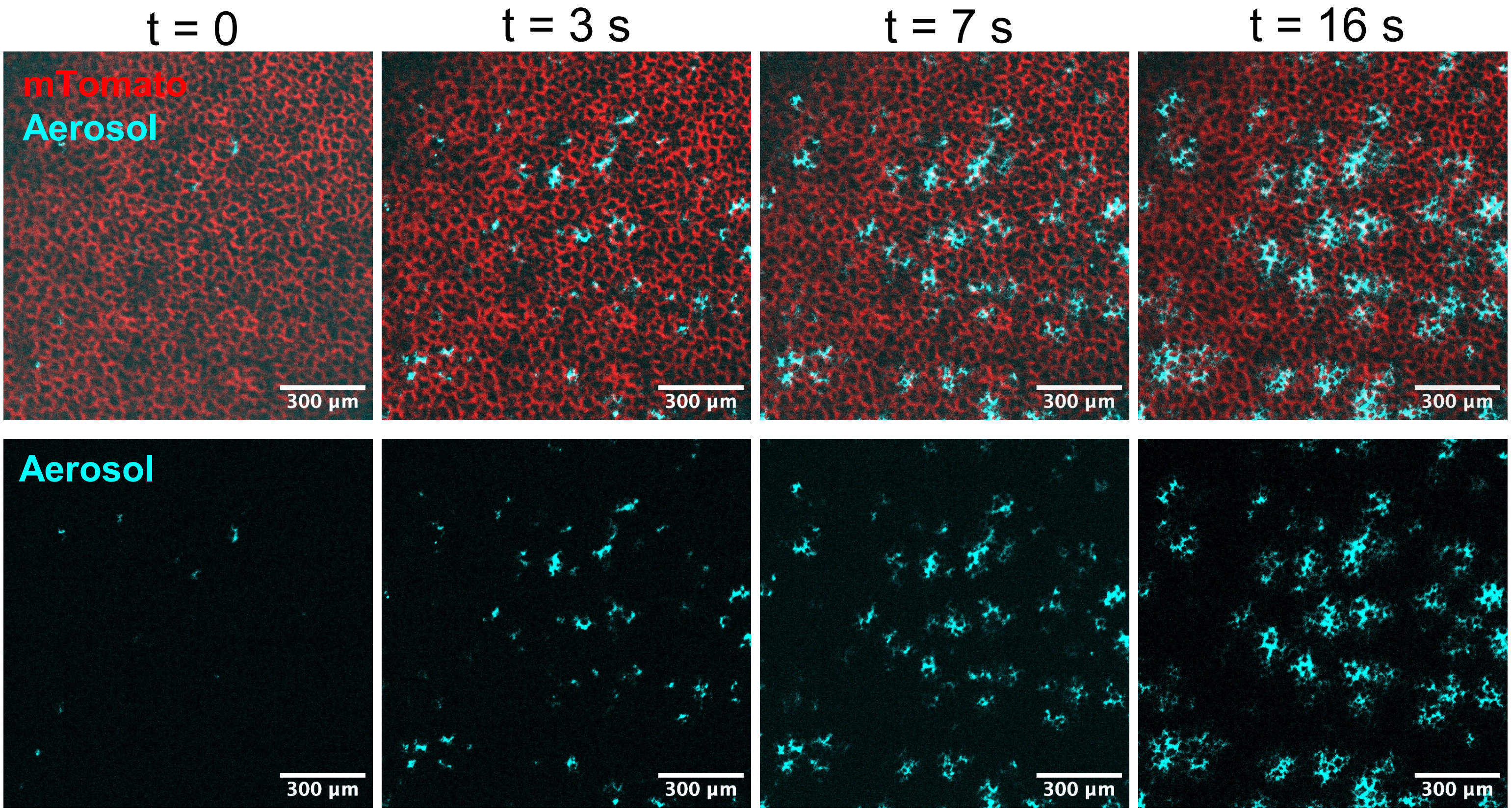


**Extended Data Figure 8 | Single droplet deposition leads to mosaic pattern formation.** Dynamic imaging of mosaic pattern formation shows that single droplet deposition over several seconds leads to the formation of mosaic pattern ”tile” structure, where droplets accumulate only in specific alveoli (tiles) while band alveoli receive zero deposition. After depositing, single droplets deform and spread along the air-liquid interface but remain confined to a small group of local alveoli.

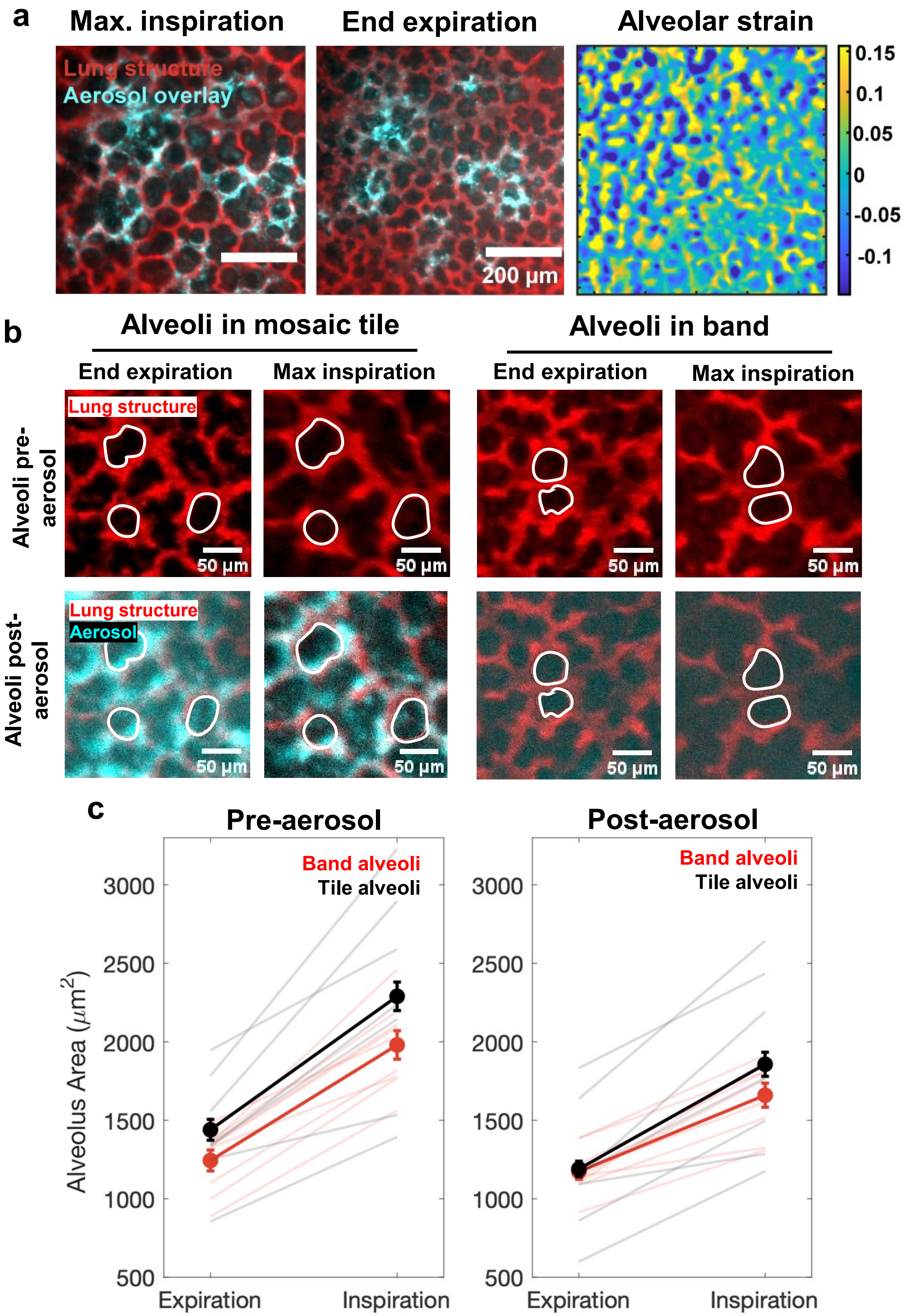


**Extended Data Figure 9 | Alveolar strain is consistent between mosaic tiles versus bands before and after aerosol delivery** (**a**) Alveoli located in tiles have similar 2D area strain to alveoli in the bands. (**b**) Individual alveoli have similar 2-D segmented area at the end expiration and max inspiration points in the ventilation cycle both before and after aerosol delivery. (**b**) Area of the alveolar airspace for band and tile alveoli is similar, quantified for for n=1 mice, where $n_{tile}$= 7 alveoli and $n_{band}$= 9 (mean ± SEM) at the end-expiration and max-inspiration points during ventilation.

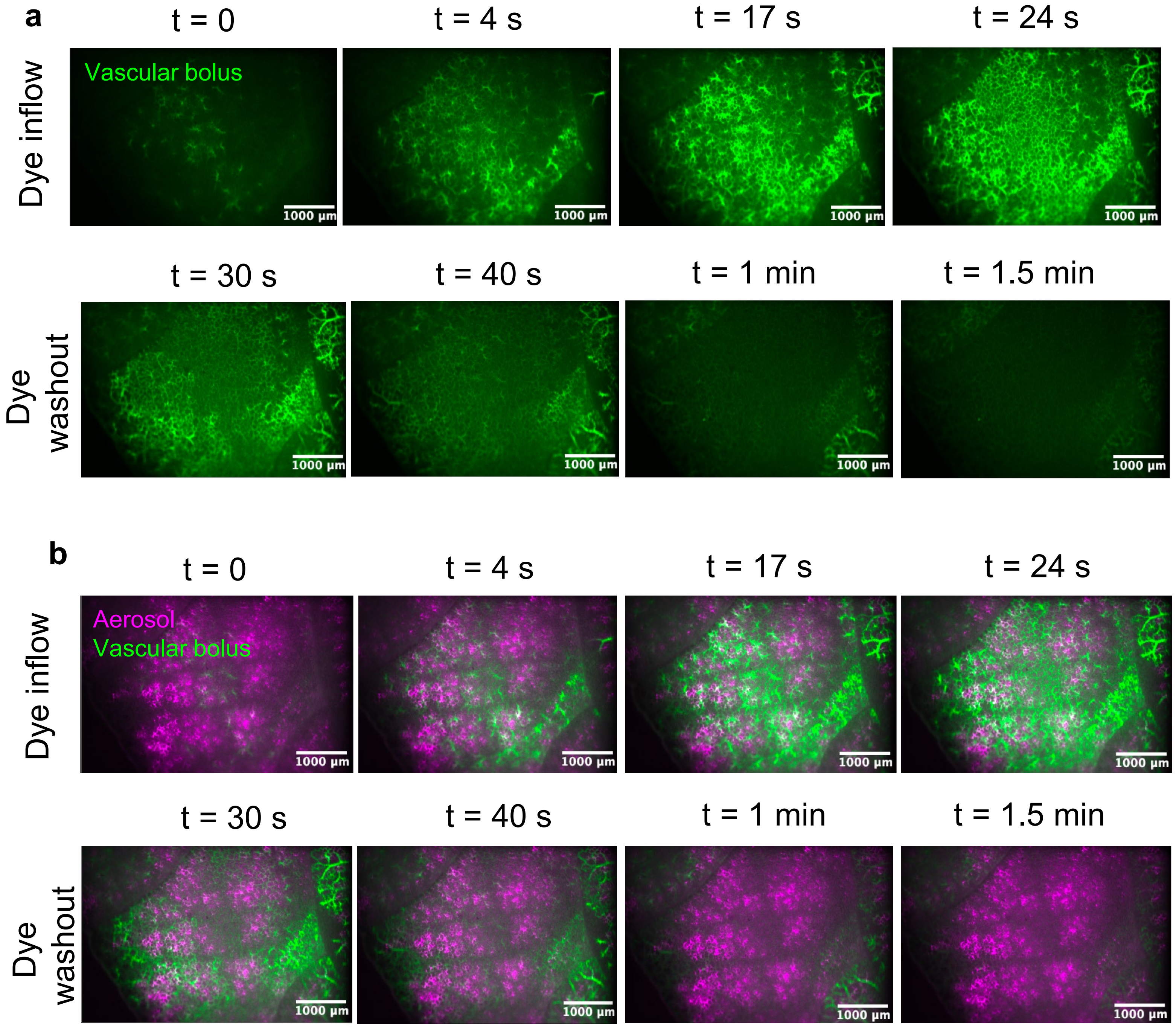


**Extended Data Figure 10 | Mosaic pattern is not spatially correlated to perfusion through pulmonary capillaries.** (**a**) Perfusion of pulmonary capillaries on the ex vivo lung surface visualized inside the crystal ribcage after administering a vascular dye bolus (50 µL, 25 mg/mL FITC albumin in saline) injected into the lung pulmonary artery during pressure-controlled perfusion (15 $cmH_2O$, ~1 mL/min flow) in the absence of ventilation. (**b**) Vascular perfusion is not spatially correlated with the mosaic pattern produced by in vivo inhalation of fluorescent aerosols prior to ex vivo perfusion (n = 1 mouse experiment, m = 2 independent ROI imaged).

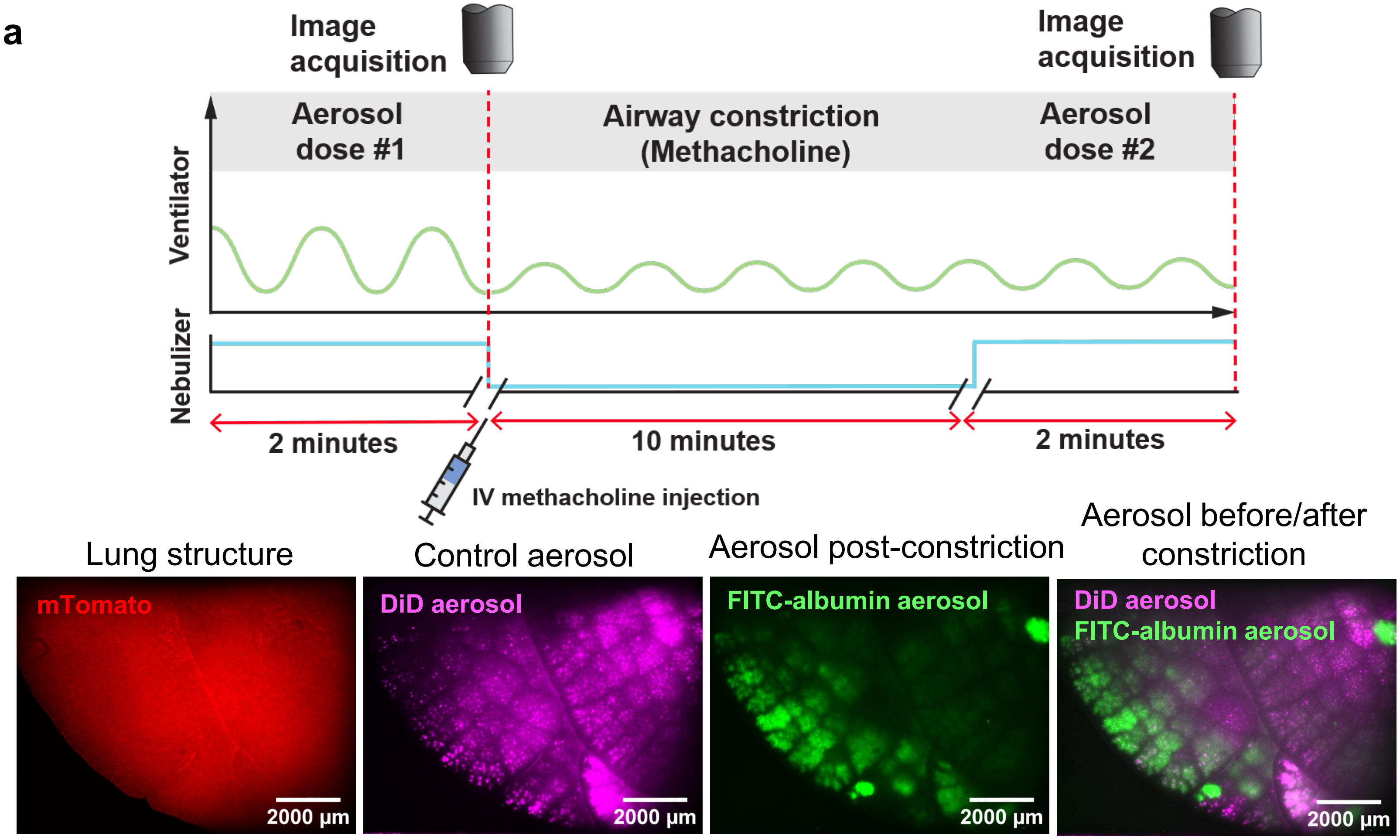


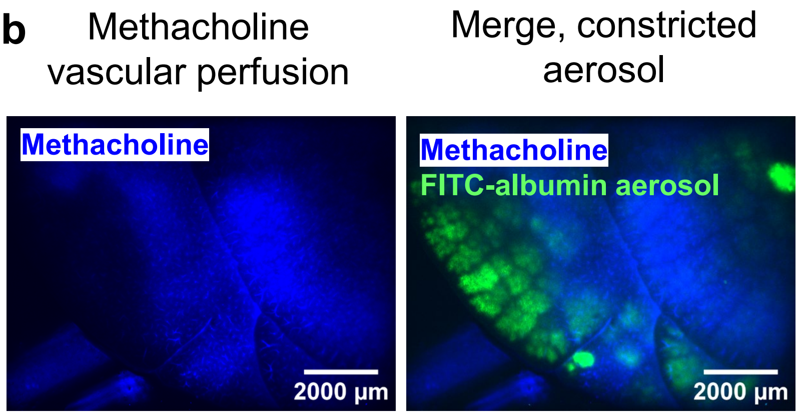


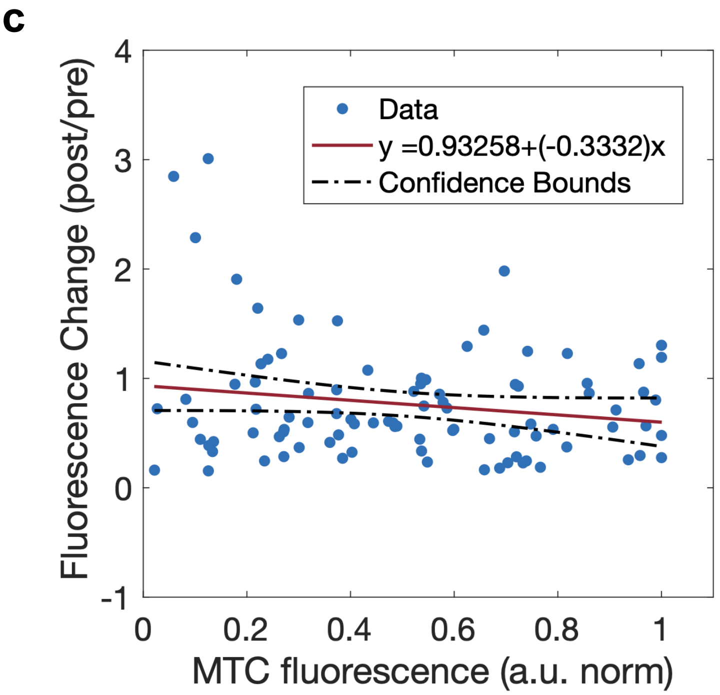


**d**

| | P-value |
|---|---|
| Y-intercept | $1.6906 \times 10^{-12}$ |
| Slope | 0.094 |

**Extended Data Figure 11 | Mosaic pattern is disrupted during airway.** (**a**) Experimental design of lung airway constriction using methacholine to directly probe the mosaic pattern as a function of airway functionality. **(b)** Airway constriction after intravascular methacholine challenge (co-delivered with cascade blue dextran, 10 kDa) causes disruption of the mosaic pattern in comparison to a control aerosol dose delivered prior to constriction, n = 4 independent lung experiments. (**c**) The distribution of methacholine on the lung surface trended towards anti-correlation with the mosaic pattern after airway constriction. Dot = individual mosaic tiles defined by control mosaic pattern, red line = linear model of relationship between methacholine perfusion signal and fluorescence fold change before and after airway constriction, dotted line = 95% confidence boundary, (**d**) table of p-values for the linear model in (c).

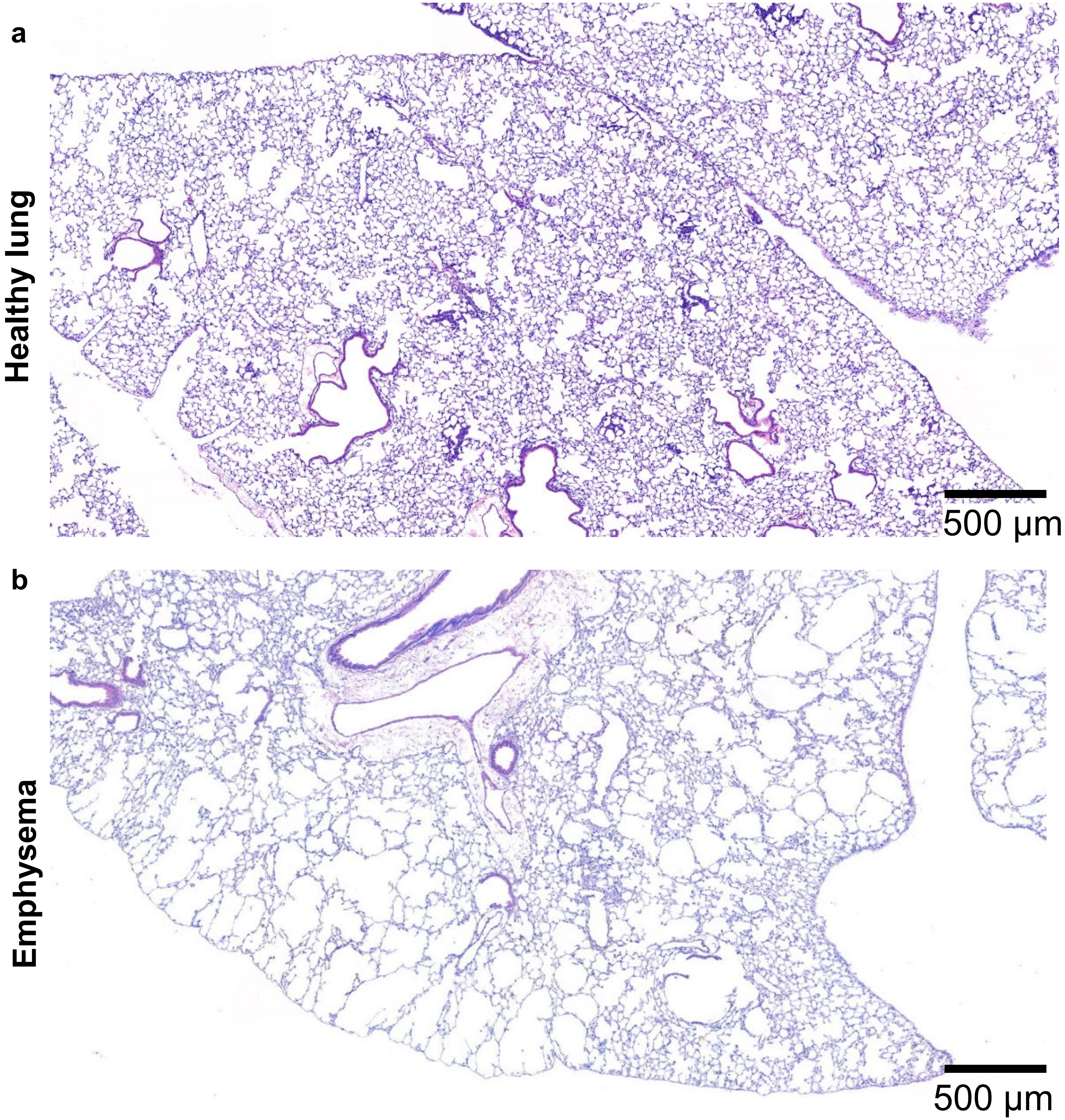


**Extended Data Figure 12 | Histology of elastase model of emphysema in mice.** (**a**) Representative hematoxylin and eosin staining of healthy mouse lung tissue (right lung), n = 3, (**b**) representative hematoxylin and eosin staining staining of mouse lung tissue after elastase injury, where large alveoli are present throughout the parenchyma due to septal wall rupture, n = 2 independent mouse experiments.

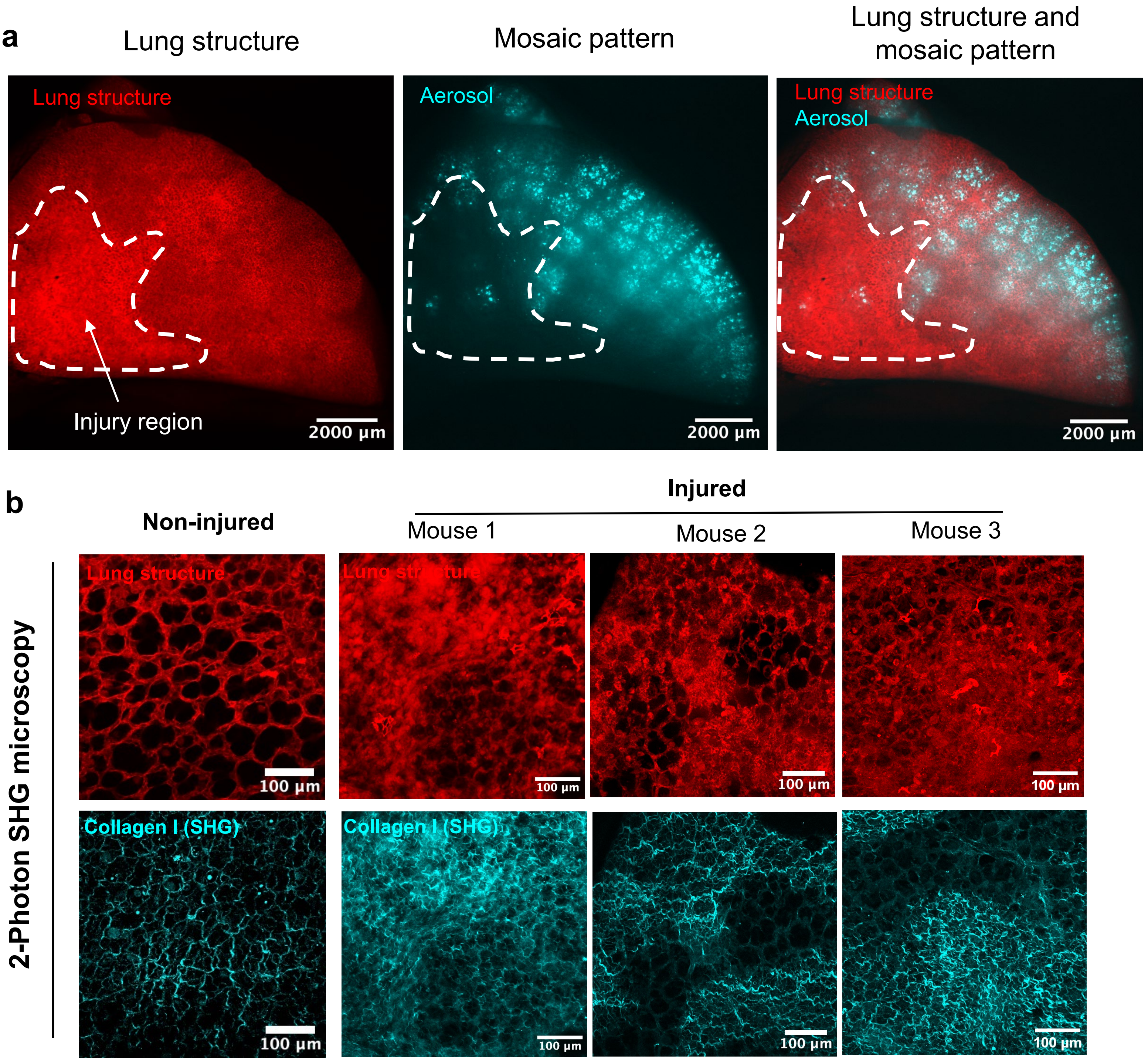


**Extended Data Figure 13 | Pulmonary fibrosis prevents formation of mosaic pattern in injured regions.** (**a**) In a mouse model of pulmonary fibrosis, the mosaic pattern tiles do not form in heavily injured regions after aerosol inhalation of cell-membrane adhesive dyes but do form in non-injured regions of the lung. (**b**) 2-Photon second harmonic generation (SHG) imaging revealed increased collagen deposition in regions heavily remodeled by bleomycin injury compared to non-injured regions.

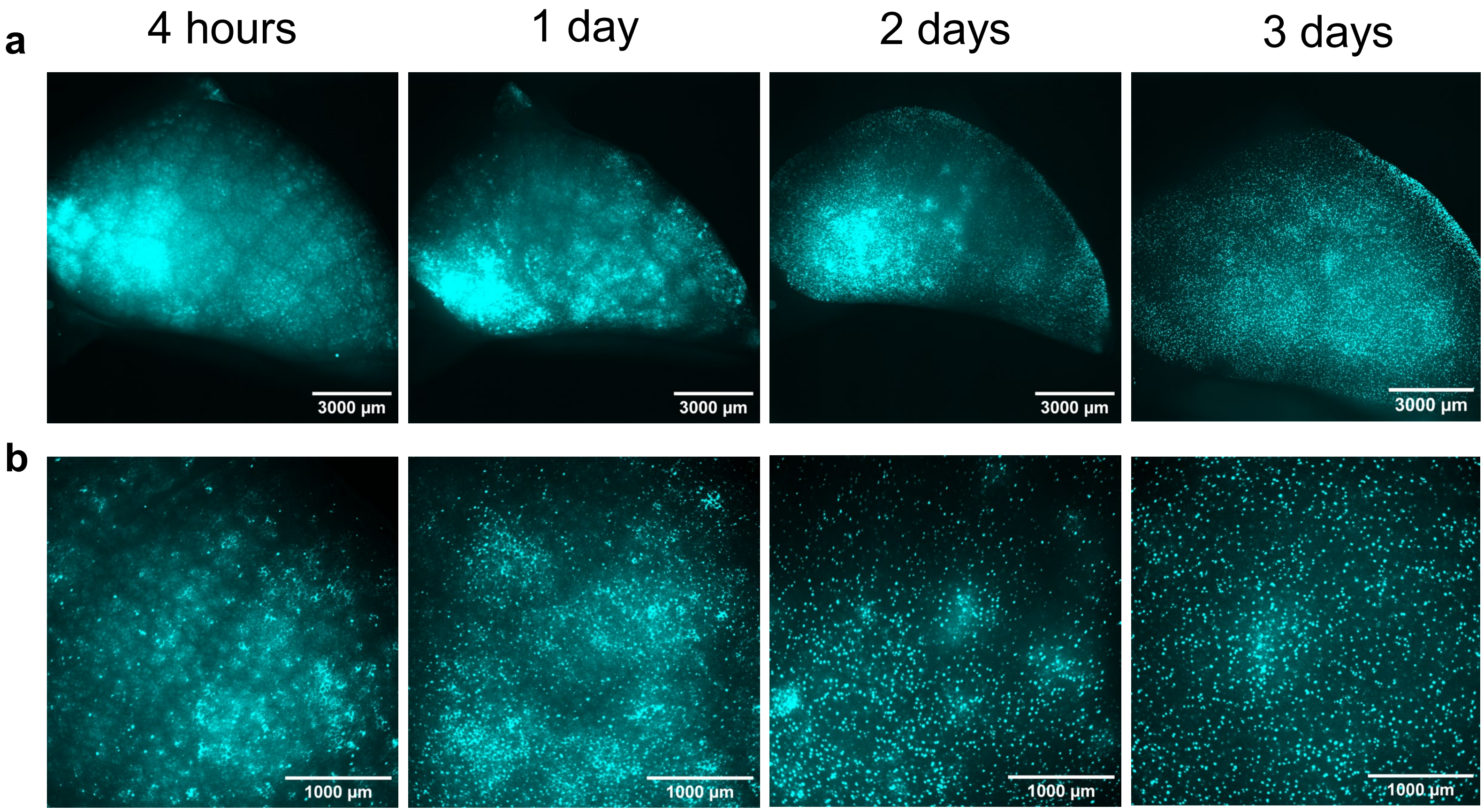


**Extended Data Figure 14 | Mosaic pattern stability over multiple days in vivo.** The mosaic pattern structure is present after administering cell-membrane adhesive aerosols in vivo via positive pressure ventilation, and sacrificing mice at 4 hours, 1 day, 2 days, and 3 days post delivery for imaging. The mosaic pattern is visible for up to two days after delivery, after which the mosaic pattern dissipates at both the (**a**) whole lung and (**b**) lobar scale, n = 1 mouse per sacrificial timepoint.

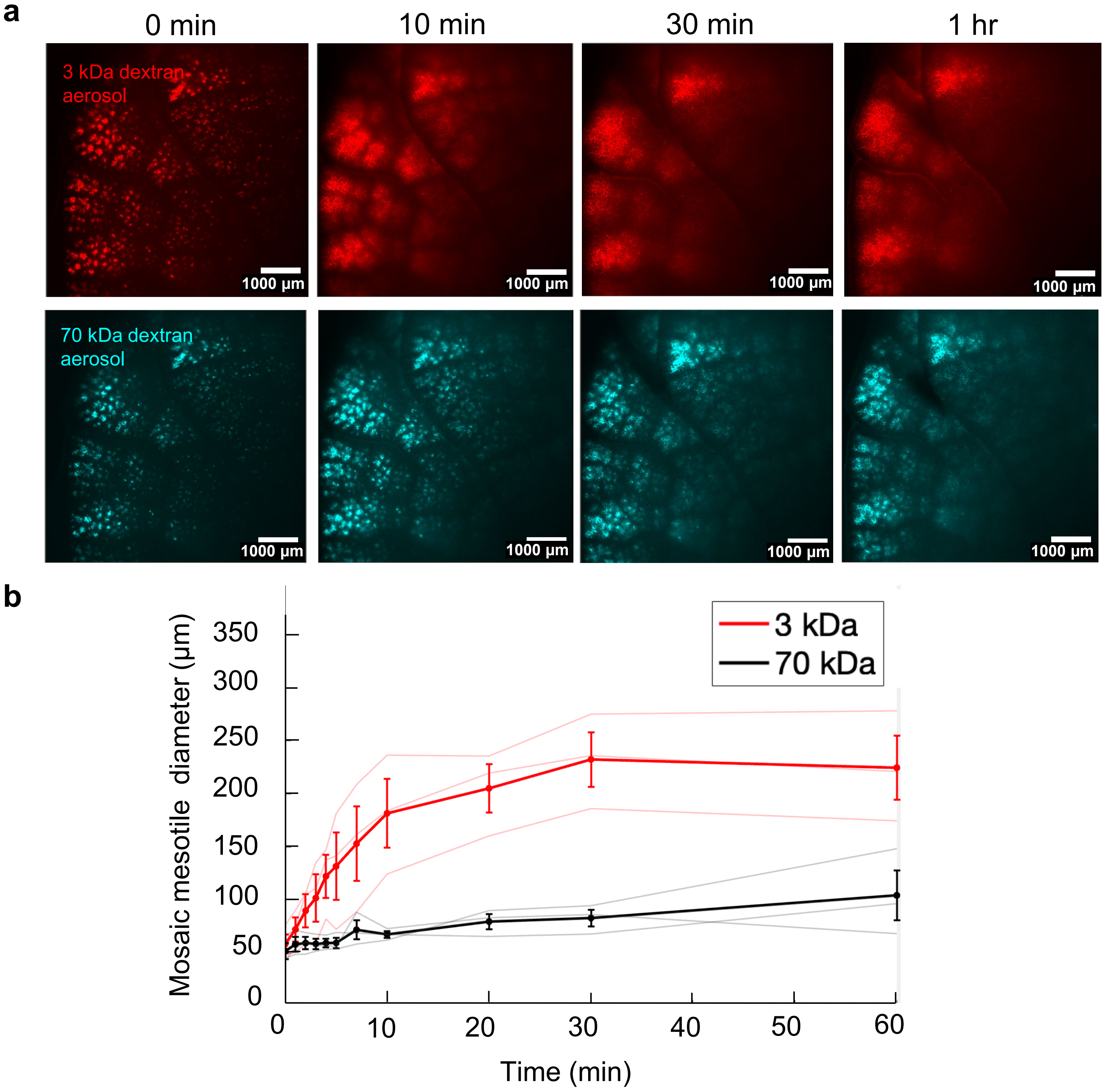


**Extended Data Figure 15 | Molecular weight aerosol redistribution is conserved across materials. (a)** Aerosol redistribution in young adult (2-6 month) for 3 kDa Texas Red-dextran co-delivered with 70 kDa FITC-dextran in saline aerosols after ex vivo inhalation under negative pressure ventilation, n = 3 mouse experiments per age group, (**b**) width of mesotiles measured over time for each fluorophore, n = 3 mice, m = 19 total ROI, (mean ± SEM).

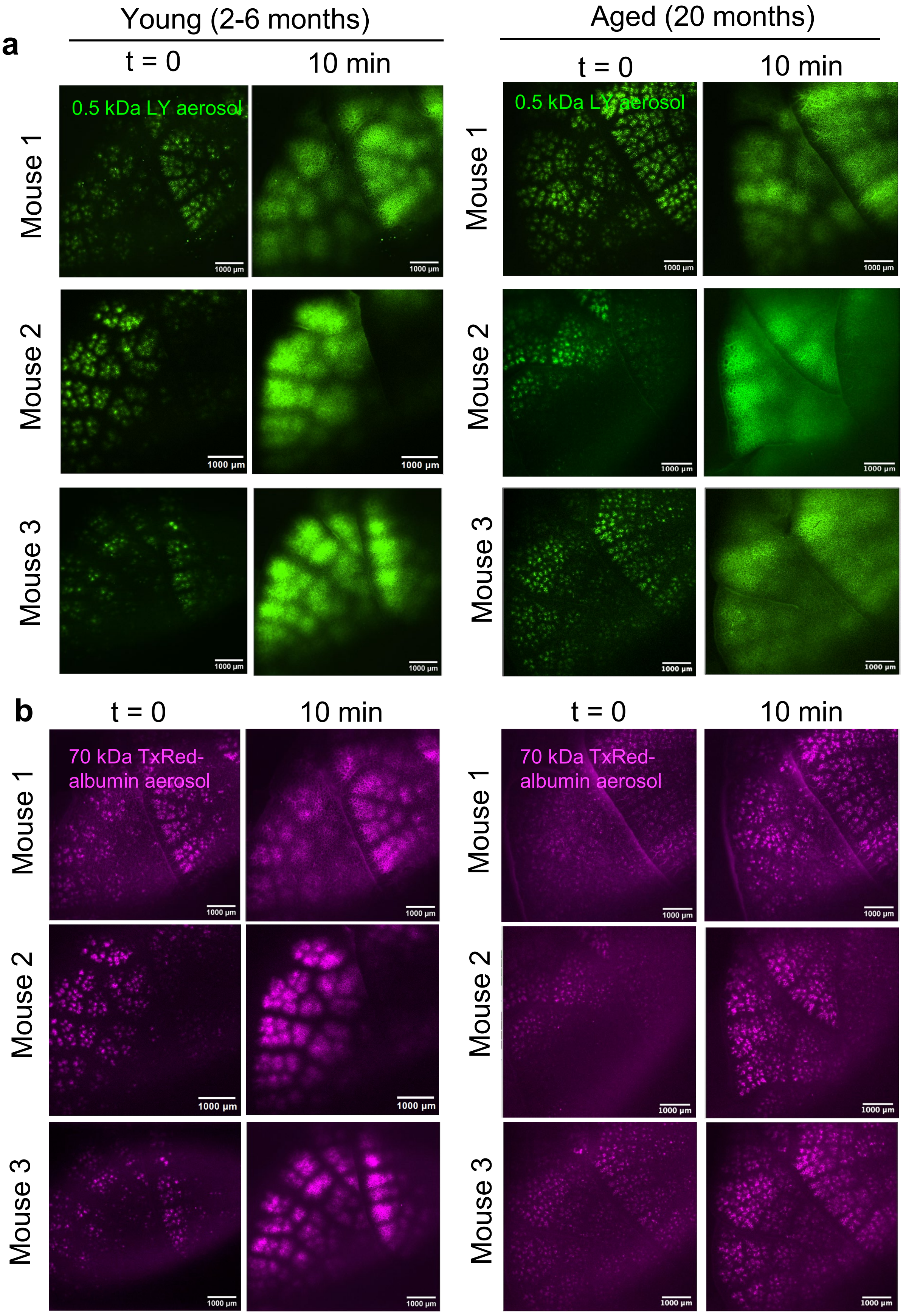


**Extended Data Figure 16 | Post deposition aerosol redistribution is molecular weight and age dependent, all replicates.** n = 3 mouse replicates for young (2-6 month) and aged (20-28 month) mouse age condition, redistribution of (**a**) 0.5 kDa Lucifer Yellow (LY) fluorescent dye co-delivered with (**b**) 70 kDa Texas Red (TxRed) albumin fluorescent dye in saline aerosols after ex vivo inhalation under negative pressure ventilation.

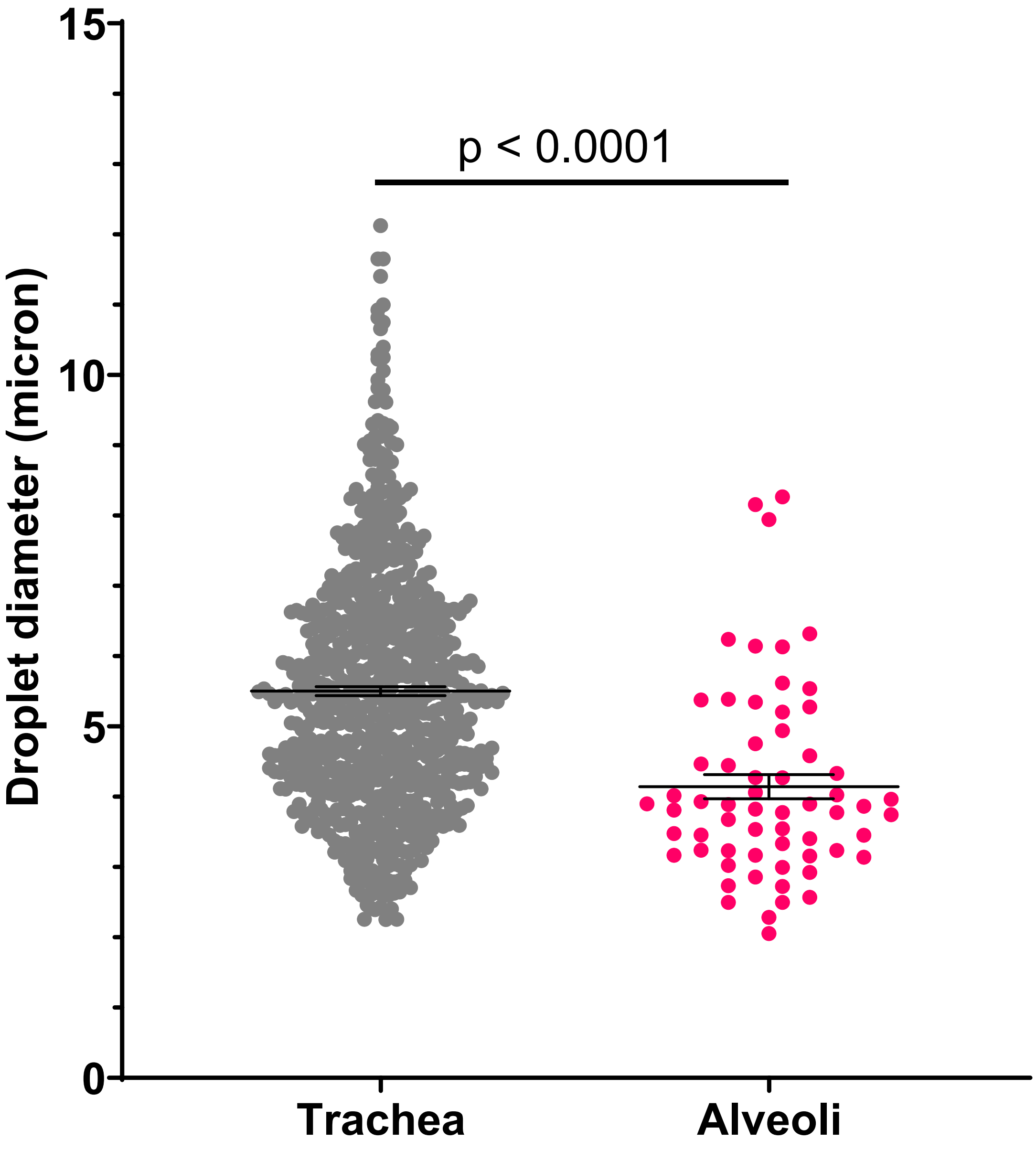


**Extended Data Figure 17 | Comparison of single droplet size distribution measured by optical microscopy via a trachea window versus in lung alveoli. (a)** Size of droplets measured for 0.5 kDa Lucifer Yellow (LY) fluorescent dye liquid droplets dissolved in saline, delivered to lungs inside crystal ribcage during ex vivo inhalation under negative pressure ventilation. Droplets were imaged using spinning disk confocal microscopy at image acquisition rate of 200 FPS either through the transparent trachea chip, or inside alveoli through the crystal ribcage, n = 2 independent mouse experiments, alveoli = 62 droplets measured, trachea = 727 droplets measured, mean ± SEM across all droplets. P-value calculated by two-tailed t-test with Welch's correction.

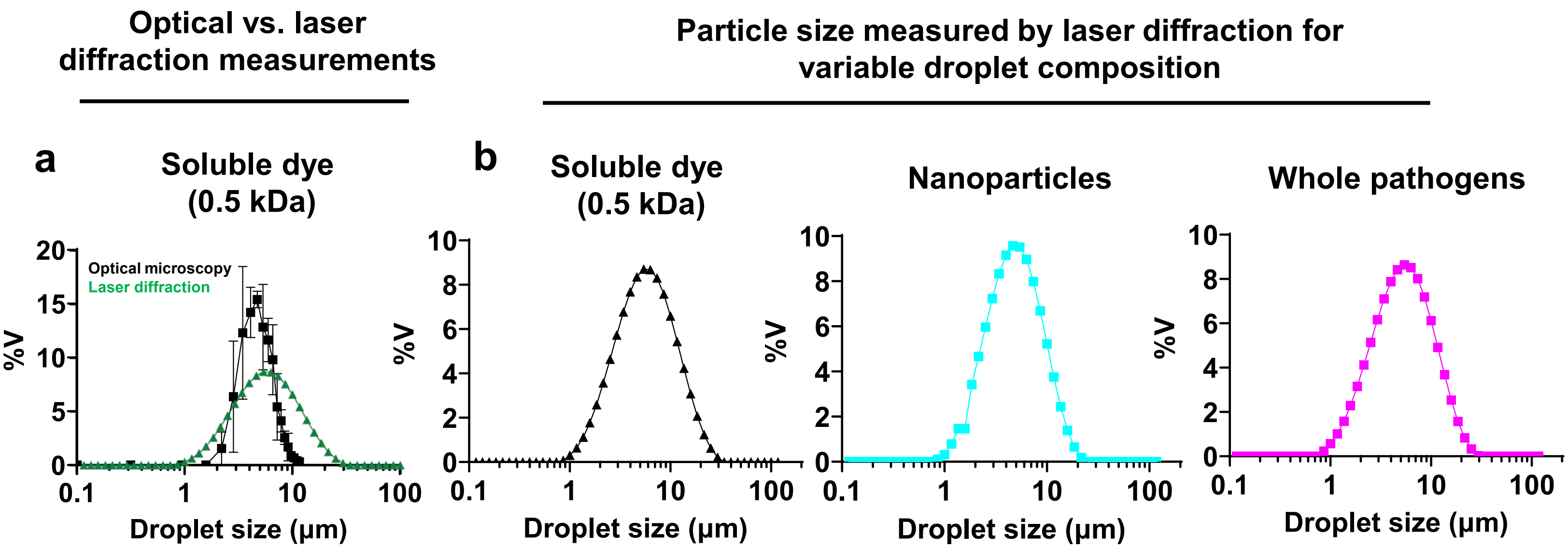


**Extended Data Figure 18 | Optical microscopy measurements are comparable to measurements obtained by a particle size analyzer for multiple materials.** (**a**) Single droplets size distribution measured using optical confocal microscopy data during real time delivery to mouse lungs inside the crystal ribcage (n = 2 independent mouse experiments), versus by laser diffraction using a particle size analyzer. (**b**) Aerosol size distributions for liquid droplets containing diverse materials are similar, measured by laser diffraction.

**a**

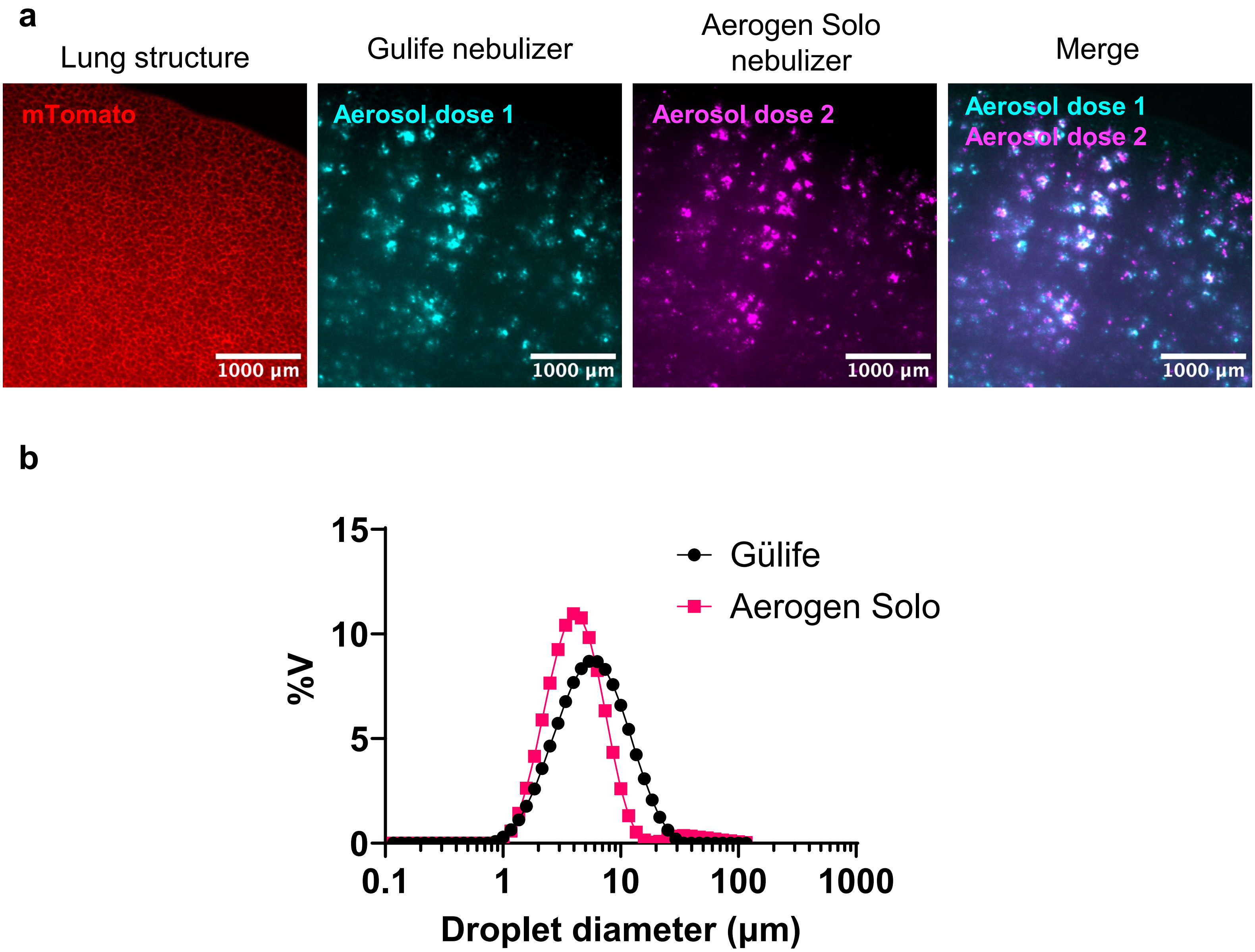


**Extended Data Figure 19 | Mosaic pattern formation is similar across dose administration using different commercial liquid nebulizers. (a)** The mosaic pattern is conserved after delivering multiple doses of liquid aerosols into the same lung using either Gülife ultrasonic mesh nebulizer and the Aerogen® Solo commercial nebulizer. (**b**) Size distribution of droplets produced by the Gülife ultrasonic mesh nebulizer and the Aerogen® Solo commercial nebulizer measured by laser diffraction using a particle size analyzer.

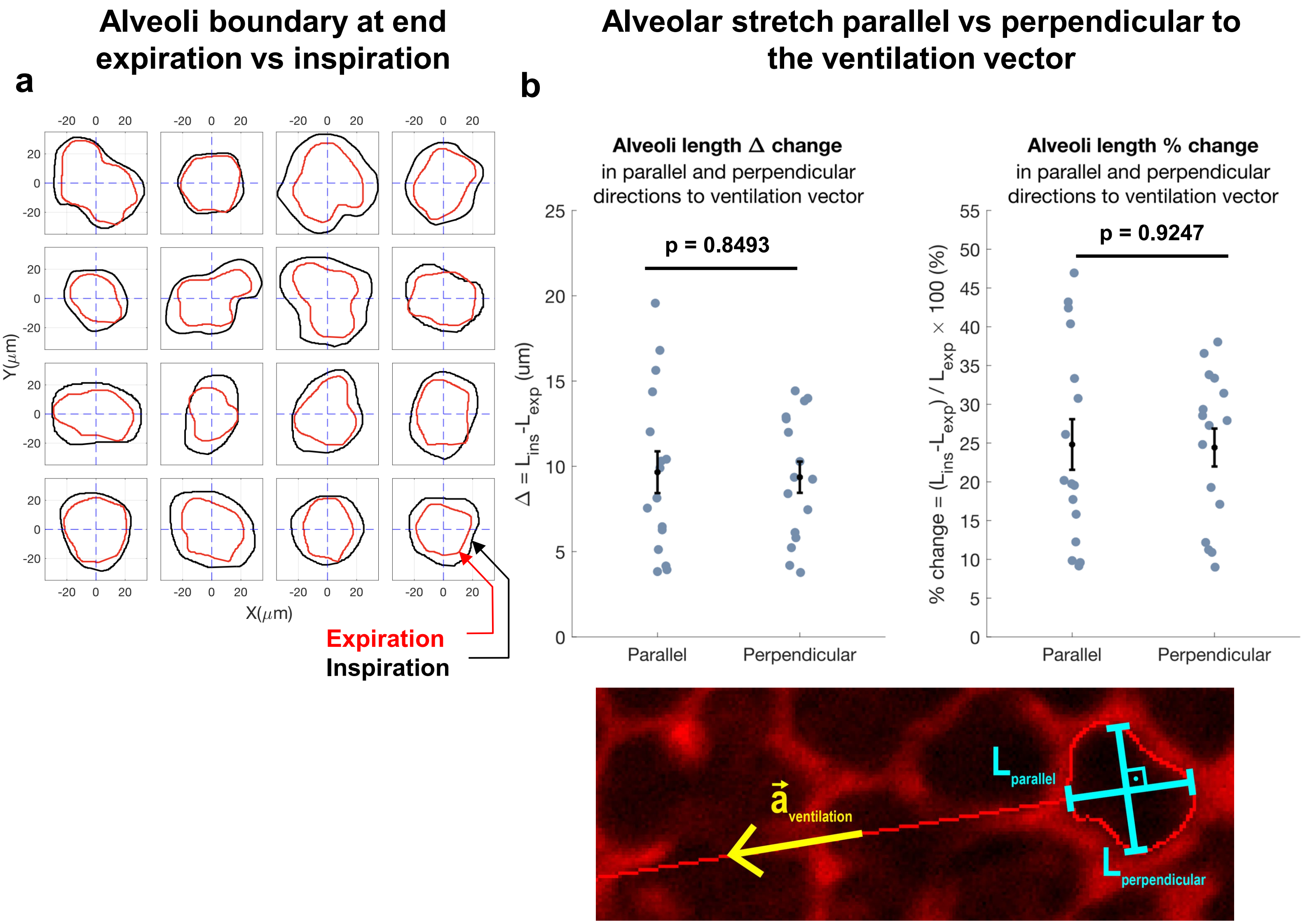


**Extended Data Figure 20 | Isotropic alveolar expansion during ventilation.** (**a**) Alveolar airspace outline manually traced at end expiration and peak inspiration during negative ventilation, where alveolar stretch between end expiration and peak inspiration is quantified as the (**b**) length change of the alveolus along the direction of the ventilation vector or perpendicular to the ventilation vector or the (**c**) percent change along or perpendicular to the ventilation vector. The ventilation vector was defined by drawing a line between the centroid of alveolus from its location at end expiration to peak inspiration. Length change or percent change is similar between parallel and perpendicular directions, indicating isometric alveolar expansion; n = 1 independent mouse experiment, m = 16 alveoli quantified (mean ± SEM), p-value calculated by paired t-test.
